# Intrinsically disordered regions in the transcription factor MYC:MAX modulate DNA binding via intramolecular interactions

**DOI:** 10.1101/2023.06.21.545551

**Authors:** Stefan Schütz, Christian Bergsdorf, Sandra Hänni-Holzinger, Andreas Lingel, Martin Renatus, Alvar D. Gossert, Wolfgang Jahnke

## Abstract

The basic helix-loop-helix leucine zipper (bHLH-LZ) transcription factor (TF) MYC is in large parts an intrinsically disordered oncoprotein. In complex with its obligate heterodimerization partner MAX, MYC preferentially binds E-Box DNA sequences (CANNTG). At promotors containing these sequence motifs, MYC controls fundamental cellular processes such as cell cycle progression, metabolism, and apoptosis. A vast network of proteins controls MYC function via intermolecular interactions. In this work, we establish another layer of MYC regulation by intramolecular interactions. We use Nuclear Magnetic Resonance (NMR) spectroscopy to identify and map multiple binding sites for the C-terminal MYC:MAX DNA binding domain (DBD) on the intrinsically disordered regions (IDRs) in the MYC N-terminus. We find that these binding events in *trans* are driven by electrostatic attraction, that they have distinct affinities, and that they are competitive with DNA binding. Thereby, we observe the strongest effects for the N-terminal MYC box 0 (Mb0), a conserved motif involved in MYC transactivation and target gene induction. We prepared recombinant full-length MYC:MAX complex and demonstrate that the interactions identified in this work are also relevant in *cis*, i.e. as intramolecular interactions. These findings are supported by Surface Plasmon Resonance (SPR) experiments, which revealed that intramolecular IDR:DBD interactions in MYC decelerate the association of MYC:MAX complexes to DNA. Our work offers new insights how bHLH-LZ TFs are regulated by intramolecular interactions, which opens up new possibilities for drug discovery.

## Introduction

The oncoprotein MYC is upregulated in more than 70 % of human cancers, and despite decades of research there is no approved MYC-targeting treatment to date^1–3^. Only one candidate, the designed MYC variant Omomyc (Omo-103), proved successful in phase I/II clinical trials so far^4^. Due to its central role in many fundamental biological processes^5^ such as cell growth and proliferation^6,7^, tumorigenesis^8–10^ and apoptosis^11,12^, MYC needs to be tightly controlled on all possible levels from gene to protein^13–18^, which involves a tight network of interacting partner proteins^19–21^.

One reason why MYC is such a difficult-to-drug target is its intrinsically disordered nature^22,23^. Scattered across the N-terminal 350 amino acid-long disordered region are short conserved sequence elements, so-called MYC boxes (Mb), that serve as binding sites for transactivating factors and other interaction partners^24–28^. While these MYC boxes appear disordered in the unbound form, they might adopt a defined structure in complex with a binding partner^29^. The C-terminus of MYC harbors the basic region and the helix-loop-helix motif (bHLH) followed by a leucine zipper (LZ), the latter promoting heterodimerization with other proteins of the bHLH-LZ family^30^. As a dimer, the bHLH-LZ motif forms the DNA-binding domain (DBD)^31^ by adopting a stable α-helical fold even in the absence of DNA. The best characterized dimerization partner of MYC is the MYC-associated factor X (MAX), a 160 amino acid protein with a bHLH-LZ motif in its central region (residues 22-102) that is flanked by disordered N- and C-terminal tails^32,33^. In contrast to MAX, which can form homodimers, MYC strictly requires heterodimerization for its DNA-binding and transcriptional activity^34–38^. MYC:MAX heterodimers preferentially bind DNA sequences that contain Enhancer box (E-Box) elements with the CACGTG hexamer as a consensus motif ^39,40^. However, promiscuous binding to E-box sequences with alterations in the core hexamer has been reported as well, especially in the case of malignant MYC upregulation^41–44^. While many drug discovery efforts target the MYC:MAX DBD, the high positive net charge and the shallow, poorly defined pocket between the two basic helices might impose a difficulty to directly target MYC at this particular site with low molecular weight compounds^1,3^.

In recent years, it became evident that intrinsically disordered regions (IDRs) are important for proper transcription factor (TF) function and regulation^45^. However, whether the IDRs in MYC have a regulatory role in addition to binding factors involved in MYC transactivation or turnover, is an aspect that has so far not been extensively explored. In this work, we were able to produce full-length MYC:MAX protein in milligram scale, wich enabled Nuclear Magnetic Resonance (NMR) spectroscopy studies to identify and map interactions between MYC IDRs, as well as between Mb’s and the MYC:MAX DBD. We used Surface Plasmon Resonance (SPR) experiments to show that conserved Mb’s, especially Mb0, are key determinants for the reduced DNA affinity of full-length MYC:MAX compared to the isolated DBD, as we find them to decelerate DNA association. SPR measurements further reveal a ten-fold higher affinity of full-length MYC:MAX to E-Box than to mutant E-Box DNA. We show that Mb’s engage the MYC:MAX DBD via a DNA-competitive mechanism, an apparently conserved mode-of-action to fine-tune DNA binding proteins^46–49^. Our work provides first evidence for direct, intramolecular Mb:DBD interactions within MYC. We find that these interactions play an important role in the regulation of MYC activity. Our results thus open up new paths to drug discovery for a direct MYC-targeting approach by utilizing the herein indentified protein-protein interactions.

## Materials and Methods

### Protein expression

All genes were codon optimized for expression in *E.coli* and cloned into modified pET vectors. MYC (UniProt P01106) 1–88, 89–150, 151–203, 203–255, 256–299, 303–351, 151–255 and 256–351 fragments carry an N-terminal His6-ZZ-TEV tag. Full-length MYC(1–439) and MYC(1– 439_C70A, C117A, C133A, C171A, C188A, C208, C300A, C342A, C438A), short MYC(1–439)_C0,C25, were cloned without any purification tag. MYC(11–34), corresponding to the Mb0 peptide, carries an N-terminal His6-MBP-TEV tag. Full-length MAX (UniProt P612244), MAX(2–160), carried an N-terminal TEV-cleavable Strep-tag. MYC:MAX DBD complexes were cloned with an uncleavable His6-tag on MYC. Mb0-MYC:MAX DBD, MbIIIa-MYC:MAX DBD and MbIIIb-MYC:MAX DBD constructs carried a TEV-cleavable His6-tag. Molecular cloning and site-directed mutagenesis were performed with an adapted strategy based on the Golden Gate Assembly method using Type IIS restriction enzymes^50^. See Table S2 for a comprehensive list of protein constructs used in this study.

*Escherichia coli* BL21 (DE3) cells (#C2527, NEB) were transformed with the appropriate plasmid and grown at 37°C to an OD_600_ of 0.8 - 1.0 in MOPS-buffered Terrific broth (TB). Overexpression of soluble proteins was induced with 0.2 - 0.5 mM IPTG at 20°C. After 15h, cells were harvested by centrifugation. Overexpression of full-length MYC into inclusion bodies was induced with 1 mM IPTG and cells were maintained at 37 °C for 5h before harvest. Proteins with stable isotopic labeling for NMR studies were overexpressed in M9 minimal medium that was supplemented with 1 g/L ^15^NH_4_Cl as the sole nitrogen source and 4 g/L D-glucose or 2 g/L ^13^C_6_-glucose as the sole carbon source. For labeling of the MYC:MAX DBD, the M9 medium was based on D_2_O.

### Purification of soluble proteins

Cells were resuspended in buffer A (50 mM Tris, pH 8, 300 mM NaCl, 1 mM DTT) complemented with 10 mM imidazole, lysozyme, 0.1 % (v/v) Triton X-100, 5 mM MgCl_2_ and 25 U/mL Turbonuclease. For ZZ- or MBP-tagged MYC constructs, cell lysis was performed by sonication on ice, while for cells containing full-length MYC, MYC:MAX complexes or MAX(2– 160) a high-pressure homogenizer was used. The lysate was cleared by centrifugation.

For His-tagged MYC proteins, the supernatant was loaded on 5 mL pre-packed Ni-NTA columns. The resin was washed with 20 CV (column volumes) buffer A mixed with 1 % buffer B (as buffer A, supplemented with 1 M imidazole). The resin-bound protein was eluted with a gradient of 5-30 % buffer B over 20 CV, followed by a gradient from 30-100% buffer B over 4 CV. Eluted proteins with cleavable His-tag were supplemented with TEV protease and dialyzed at 4 °C overnight against buffer C (20 mM HEPES, pH 7.3, 100 mM NaCl, 2 mM DTT). Dialysates were cleared from precipitate by filtration and applied to 5 mL Ni-NTA resin in a gravity flow column equilibrated with buffer D (20 mM HEPES, pH 7.3, 500 mM NaCl, 1 mM DTT) supplemented with 10 mM imidazole. The column was washed with 3 CV buffer D supplemented with 20 mM imidazole. The resin-bound purification tag was eluted with 5 CV buffer B. Fractions containing the cleaved target protein were further purified by size exclusion chromatography (SEC). SEC was performed on HiLoad 16/600 Superdex 75 columns (Cytiva) in buffer E100 (20 mM HEPES, pH 7, 100 mM NaCl, 2 mM TCEP).

For MYC:MAX DBD complexes, buffers A and B were additionally supplemented with 10 % (v/v) glycerol. After elution from the IMAC resin, the NaCl concentration was reduced to 200 mM by dilution with 0.5 Vol H_2_O. The diluted complexes were further purified on a 5 mL HiTrap Heparin column (Cytiva) equilibrated in buffer F (25 mM Tris, pH 8, 200 mM NaCl). The target proteins were separated from DNA and contaminants via a 10 CV wash step with 10 % buffer G (25 mM Tris, pH 8, 2 M NaCl), followed by a gradient from 10-50 % buffer G over 40 CV and 50-100 % buffer G over 4 CV. Fractions containing MYC:MAX complexes were pooled and purified to homogeneity via SEC as above. MAX(2–160) was purified as described before^51^.

For the Mb0 peptide that carries a TEV-cleavable MBP-tag the dialysis step after elution from the IMAC resin was omitted. The TEV-cleaved peptide was directly subjected to SEC (HiLoad 16/600 Superdex 30, Cytiva) in 50 mM NH_4_HCO_3_. Fractions containing the target peptide were pooled, flash-frozen in liquid nitrogen and lyophilized. Dry peptide was dissolved in 100% d_6_-DMSO to a final concentration of 10 or 20 mM.

### Purification of full-length MYC from inclusion bodies

Cells from 2 L culture were resuspended in 50 mL cold IB buffer (50 mM Tris, pH 8, 5 mM EDTA, 5 mM Benzamidine, 1 mM DTT) complemented with lysozyme, 5 mM MgCl_2_, 25 U/mL Turbonuclease and complete protease inhibitor cocktail (Roche). After cell lysis, the inclusion body pellet was washed with 30 mL IB buffer followed by a wash with 30 mL water. Afterwards, the pellets were dissolved in buffer U7 (7 M Urea, 25 mM HEPES, pH 7.6, 300 mM NaCl, 15 % v/v glycerol, 1 mM DTT, 0.1 mM EDTA) supplemented with complete protease inhibitor cocktail (Roche) to a protein concentration of about 5 mg/mL. Any insoluble debris were removed by centrifugation. The supernatant was flash-frozen in aliquots and stored at -80 °C. Typically, at this step more than 50 mg protein were obtained from 1 L expression medium. LC-MS analysis confirmed cleavage of the N-terminal methionine stemming from the translation start site, resulting in MYC(2–439).

### Refolding of full-length MYC:MAX

Full-length MYC:MAX complexes were reconstituted by diluting equimolar amounts of MYC(2–439) and MAX(2–160) into buffer U6 (as U7, but with 6M urea) to a final concentration of 0.25-0.3 mg/mL. We typically refold 25-30 mg MYC with an appropriate amount of MAX, corresponding to not more than 0.5 L expression volume for each of the two proteins. MYC:MAX complexes were refolded by rapid, stepwise dialysis. Dilute MYC:MAX solution in buffer U6 was transferred to a dialysis bag with 12-14 kDa MWCO (SpectraPor) and dialyzed for 1 h at 20 °C against 2 L buffer U3 (as U7, but with 3 M urea), followed by dialysis for 1 h at 20 °C against 2 L buffer U1 (as U7, but with 1 M urea), for 1 h at 20 °C against 3 L buffer U.5 (as U7, but with 0.5 M urea), and finally overnight at 4 °C against 4 L buffer U0 (as U7, but without urea). Refolded MYC:MAX was further purified via ion exchange on a pre-packed 5 mL HiTrap Heparin column (Cytiva) equilibrated in buffer F. The column was washed with 5 CV buffer F and MYC:MAX was eluted with a steep gradient from 0 to 100% buffer G over 10 CV, followed by 5 CV 100% buffer G. Fractions containing full-length MYC:MAX were pooled and concentrated in Amicon Ultra-15 centrifugal filters with 10 kDa MWCO (Millipore) to 3 mL. Final purification to homogeneity was achieved on a HiLoad 16/600 S75 column (Cytiva) in buffer E300 (20 mM HEPES, pH 7, 300 mM NaCl) supplemented with 2 mM TCEP. Fractions of 0.5 mL were collected and only four to five fractions at the elution peak maximum were pooled, flash-frozen and used in subsequent experiments without further concentration.

### 13C-MMTS, Iodoacetamide and TEMPO spin labeling

Labeling of refolded full-length MYC:MAX with ^13^C-MMTS (^13^C-methyl methanethiosulfonate; Sigma-Aldrich) was performed after elution from the Heparin column (see above). Purified Mb0-MYC(352–437):MAX(2–160) was labeled after exchanging the buffer to TCEP-free buffer E100. Fully reduced Mb0 peptide was prepared by diluting a 10 mM solution of MYC(11–34) in 100% d_6_-DMSO by about 100-fold in buffer E100 supplemented with 5 mM DTT, followed by buffer exchange to TCEP-free buffer E100 in a centrifugal filter (3 kDa MWCO). Proteins were incubated with a 2-fold molar excess of ^13^C-MMTS (100 mM stock in 100% d_6_-DMSO) at 4 °C overnight (ON). Acetamide-labeling was achieved by addition of a 4.5-fold molar excess of iodoacetamide (Sigma-Aldrich; 50 mM stock in 100% d_6_-DMSO) and the labeling reaction was incubated ON at 4 °C in the dark. TEMPO-labeling was performed with a 10-fold molar excess of 4-maleimido-TEMPO (4-Maleimido-2,2,6,6-tetramethyl-1-piperidinyloxy; Sigma-Aldrich; 200 mM stock in 100% d_6_-DMSO) and the peptide was incubated ON at room temperature in the dark. Unreacted label and side products were removed via SEC as described above, maintaining buffer conditions that are free of reducing agents. LC-MS confirmed the completeness of the labeling reaction without any secondary site labeling on MYC or MAX.

### Proteins, Peptides and DNA from commercial sources

MYC(151–439):MAX(2-160) was purchased from Biortus (Jiangyin, China).

MAX(2–21)_pS2,pS11 (Ac-pSDNDDIEVEpSDEEQPRFQpSA-NH_2_) was purchased from Biosyntan (Berlin, Germany). Ac-denotes an acetylated N-terminus and -NH_2_ denotes a C-terminal amide, while pS is for phospho-serine.

Duplex DNA probes for SPR and NMR measurements were purchased from IDT (Leuven, Belgium). See Table S1 for a complete list of DNA duplexes used in this study.

### NMR spectroscopy and data analysis

All NMR samples were prepared in buffer E100 or E300 and contained 5% (v/v) D_2_O. NMR spectra were recorded at 283 K for the disordered MYC fragments and at 298 K for MYC:MAX complexes on Bruker AVIII-600 or AVIII-800 spectrometers with cryogenic probe-heads. For MYC:MAX complexes, transverse-relaxation optimized spectroscopy (TROSY) techniques were employed together with deuteration to reduce relaxation^52^. Backbone and side-chain resonances were assigned using standard triple-resonance CBCANH, CBCA(CO)NH, HNCO, HN(CA)CO and (H)CC(CO)NH experiments^53^. Samples for assignment purposes contained 665 μM and 900 μM protein for MYC(151–255) and MYC(256–351), respectively. Several low-intensity resonances could be linked to *cis*-prolines, which were readily identified based on their characteristic Cβ, Cγ and C’ chemical shifts^54^.

NMR binding experiments were performed with 25 μM or 50 μM NMR-active protein and unlabeled protein or DNA as indicated in the figures. Heteronuclear single quantum correlation spectroscopy (HSQC) experiments were used to monitor changes in the amide proton and nitrogen chemical shifts. The combined chemical shift difference Δδ_comb_ of a given resonance in absence and presence of a binding partner was calculated using equation (1):

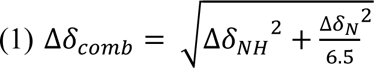

where Δδ_NH_ and Δδ_N_ are the differences in the amide proton and nitrogen chemical shift, which are weighted by a scaling factor R_scale_ = 6.5 ^55^. Significant CSPs were determined by calculating a cut-off of 2σ from the trimmed mean^56^. For intensity analysis, the ratio of peak intensities in HSQC spectra of bound (I_bound_) and free (I_free_) protein was calculated as I_bound_/I_free_.

NMR titration data was fitted globally in Origin Pro 2021 (OriginLab) using equation (2):

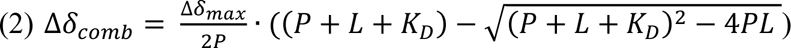

where P and L are the constant protein and variable ligand concentrations, respectively, Δδ_comb_ is the observed chemical shift difference calculated from equation (1) and Δδ_max_ is the maximal chemical shift difference reached at saturation^57^.

NMR experiments with ^13^C-MTC- and TEMPO-labeled protein were carried out in buffer E100 or E300 lacking TCEP. A MYC:MAX DBD NMR sample was prepared with an equimolar amount of TEMPO-labeled Mb0 and measured under non-reducing conditions. Subsequently, 500 μM ascorbic acid were added to the NMR sample and the measurement was repeated under reducing conditions. A control sample contained only MYC:MAX DBD and 500 μM ascorbic acid, to rule out interactions of ascorbic acid with the DBD. The ratio of peak intensities in ^1^H-^15^N TROSY spectra recorded under oxidizing (I_OX_) and reducing (I_RED_) conditions was calculated as I_OX_/I_RED._

All NMR data was processed in Topspin 3.2 or 4.1 (Bruker). CCPN 3.0 ^58^ was used for analysis and visualization of NMR data and Pymol (pymol.org) was used to produce figures of protein structures. Backbone resonance assignments for MYC(151–255) and MYC(256–351) have been deposited in the BMRB under accession codes 51636 and 51635, respectively.

### SPR

Surface plasmon resonance (SPR) experiments were performed on Biacore 8K^+^ or T200 instruments (Cytiva). Biotinylated DNA duplexes were immobilized (up to 100 RU) on a CM4 chip which has been coated with neutravidin via amine-coupling (approx. 2000 RU). Protein samples were diluted into SPR buffer (25 mM HEPES, pH 8, 150 mM NaCl, 1 mM TCEP, 0.01 % Tween-20, 3 mM EDTA). Protein solutions (typically ranging from 0.49 nM up to 1000 nM) were applied to the flow cell at a flow rate of 75 μL/min and association was monitored for 160 s. Subsequently, dissociation in SPR buffer was monitored for 190 s. For equilibrium fits, the average response signal between 150 s and 155 s association time is considered. In addition, MYC(352– 437):MAX(2–160) was allowed to associate for a total of 600 s at a flow-rate of 30 μL/min, the response signal between 235 and 240 s was considered for equilibrium fits, and dissociation was monitored for 300 s. Equilibrium K_D_ values were fitted to a 1:1 binding model using equation (3):

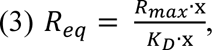

where R_eq_ is the equilibrium response signal at a given protein concentration x and R_max_ is the maximum response signal. R_eq_ values were finally normalized using R_max_ to obtain values for “fraction bound”.

## Results

### Preparation of full-length MYC:MAX in milligram scale

Structural and biophysical studies of full-length MYC have long been hampered by the inability to produce full-length MYC in sufficient quantity and quality. The majority of *in vitro* studies thus relied on shorter MYC constructs, that comprise either the folded C-terminal basic helix-loop-helix leucine zipper (bHLH-LZ) motif or portions of the N-terminal transactivation domain (TAD). In this regard, refolding of MYC:MAX bHLH-LZ dimers from the individual denatured proteins is an established method to obtain pure and homogeneous heterodimers^30,36,59^. For this study, we improved a previously published protocol^60^ to also enable milligram-scale preparations of full-length MYC:MAX complex from recombinant proteins expressed in *E. coli* (see Materials and Methods). Essentially, we were able to refold and purify homogenous full-length MYC:MAX at concentrations of up to 40 μM.

### Disordered regions in full-length MYC reduce the DNA binding affinity of the MYC:MAX TF

When we assessed DNA binding of our reconstituted full-length MYC:MAX complex, we noticed that it exhibited a remarkably lower affinity towards E-Box DNA than what has been reported by us and others for the isolated MYC:MAX DNA binding domain (DBD)^51,59,61,62^. We therefore tested three different MYC:MAX complexes, that varied in the length of the MYC protein, for their binding to a set of four different DNAs using Surface Plasmon Resonance (SPR; Figure 1). We chose the canonical E-Box sequence (CACGTG), a negative control DNA (TTAGCA) and two mutant E-Box sequences from the literature (Figure 1B)^43^. The mutant E-Box sequences carried two mutations in the core hexamer either in the middle of the recognition site (mutant E-box 1, CATATG, referred to as “low-affinity E-Box”) or at the ends (mutant E-box 2, AACGTT, referred to as “non E-Box”), which still allow for specific recognition by MYC:MAX, albeit with weaker affinity.

**Figure 1.**
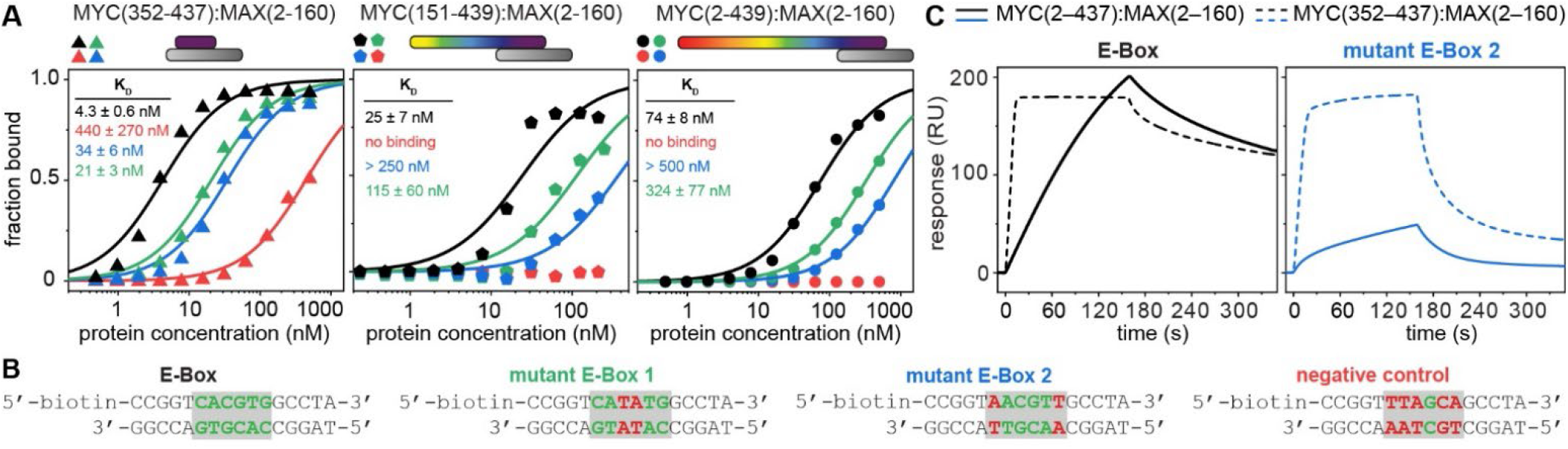
Disordered regions in MYC reduce the affinity for DNA. (A) Steady-state affinity analysis for binding of MYC(352–437):MAX(2–160) (left, triangles), MYC(151–439):MAX(2– 160) (middle, pentagons) and MYC(2-439):MAX(2–160) (right, circles) to immobilized E-Box (black), mutant E-Box 1 (green), mutant E-Box 2 (blue) or a negative control DNA (red). Values represent mean K_D_ ± SD obtained from a fit (solid lines) with equation (3), assuming a 1:1 binding model. Schematics of the MYC:MAX complexes are shown above the graphs, where MAX is in greyscale and MYC is colored with a rainbow gradient. (B) Duplex DNA oligonucleotides used in the SPR experiments. The core E-Box hexamer is in green with gray background, altered nucleotides are in red. (C) Representative SPR sensorgrams for binding of MYC(2–439):MAX(2– 160) and MYC(352–437):MAX(2–160) (solid and dotted lines, respectively) to E-Box (left, black) and mutant E-Box 2 DNA (right, blue) at 62.5 nM protein concentration, illustrating the reduced association rates for the full-length complex.

We find that all three tested MYC proteins MYC(2–439), MYC(151–439) and MYC(352–437), in complex with full-length MAX, bind strongest to E-Box DNA. The affinity towards the mutant E-Box sequences was reduced (Figure 1A), and the negative control DNA was weakly bound only by the MYC(352–437):MAX(2–160) complex. Compared to the latter, the affinity of full-length MYC:MAX is reduced by 15- to 20-fold for the canonical and mutant E-Box sequences. MYC:MAX complexes with the intermediate MYC(151–439) protein show affinities between those for MYC(352–437):MAX(2–160) and full-length MYC:MAX (Figure 1A). We thus conclude that the N-terminal disordered part of MYC weakens the intrinsic affinity of the MYC:MAX DBD toward on- and off-target DNA sequences.

### Expression strategy for disordered MYC fragments

Sequence analysis of MYC and MAX proteins suggested that the observed effects on DNA binding might be attributed to interactions between the acidic disordered TAD and the basic DBD. As these are difficult to characterize in the full-length protein, we set out to identify potential MYC:MYC interactions using a divide-and-conquer approach. Taking the established N- and C-terminal fragments MYC(1–88) and MYC(352–437) as a basis^30,63^, we divided the remaining central part into five constructs with a length of about 50 amino acids each (Figure 2A and B), while preserving the conserved MYC box (Mb) motifs. These constructs were expressed in large amounts in *E. coli* (1–20 mg per L culture) as cleavable fusion proteins. We found also the longer fragments 151–255 and 256–351 to be well-behaved with regards to expression yields, solubility and stability, which enabled us to obtain NMR resonance assignments of these two constructs (Figure S4 and S5). As the isolated MYC bHLH-LZ motif is prone to aggregation^31^, MYC(352– 437) was co-expressed with its cognate parter MAX(22–102) and purified as a heterodimer to yield highly soluble protein. Together, MYC(352–437) and MAX(22–102) form the DNA-binding domain and are devoid of long unstructured stretches (Figure S1). In the following, the heterodimer MYC(352–437):MAX(22–102) is denoted as MYC:MAX DBD.

**Figure 2.**
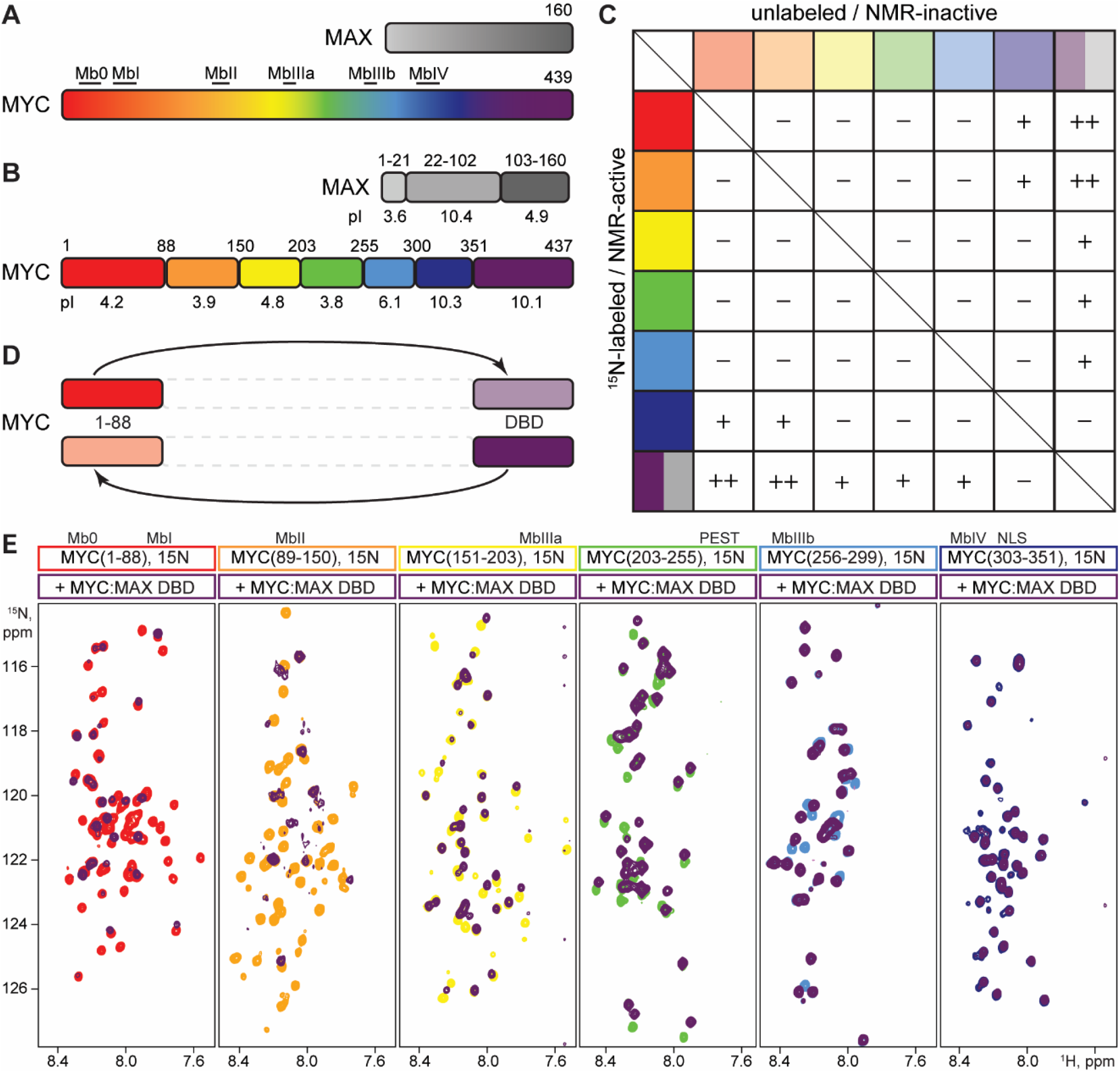
Disordered and structured regions of MYC interact with each other. (A) Schematic representation of full-length MYC and MAX. Locations of the conserved MYC Box (Mb) motifs are indicated. (B) Fragmentation of MYC to obtain well-expressing, soluble proteins. MYC(352– 437) (purple) contains the bHLH-LZ motif, which heterodimerizes with the respective motif in MAX(22–102) (grey), thereby forming the DNA binding domain. The isoelectric points and fragment boundaries are indicated. (C) Summary of NMR binding experiments and MYC:MYC interactions identified in this study. ^15^N-labeled MYC fragments (bright colors) were tested for binding to all other (unlabeled) fragments (pale colors). “+” denotes binding (presence of CSPs), “++” denotes strong binding (CSPs observed and significant decrease in signal intensities) and “– “ denotes no binding (absence of CSPs at 500 μM ligand concentration). The bipartite purple and grey field denotes the heterodimeric MYC(352–437):MAX(22–102) complex (henceforth MYC:MAX DBD). (D) NMR binding results are consistent regardless of which fragment is NMR-active. (E) ^1^H-^15^N HSQC spectra of 50 μM ^15^N-labeled MYC fragments in absence (red: MYC(1– 88), orange: MYC(89–150), yellow: MYC(151–203), green: MYC(203–255), blue: MYC(256–299), dark blue: MYC(303–351)) and in presence of a 10-fold molar excess of the MYC:MAX DBD (purple). Presence of CSPs or decrease in peak intensities indicates binding. This dataset corresponds to the last column of the matrix shown in (C).

### Detection and characterization of interactions within MYC

To identify interactions between the various MYC fragments, we used Nuclear Magnetic Resonance (NMR) spectroscopy. NMR is a particularly useful method for this kind of studies since it can robustly detect weak protein-protein interactions. NMR spectra of an isotope-labeled (and thus NMR-active) protein were recorded in absence and presence of an unlabeled potential binding partner. Upon binding, the chemical environment of residues in the binding site changes. This leads to perturbations of the chemical shifts of the affected nuclei and ultimately to altered positions of the respective peaks in the NMR spectrum. Conversely, the absence of chemical shift perturbations (CSPs) is an indication that two potential binding partners do not directly interact with each other, at least under the given assay conditions. We expressed the seven MYC fragments in an NMR-active (^15^N-labeled) form and recorded ^1^H-^15^N correlation spectra in absence and presence of each of the six other fragments (Figure 2C, and Figure S2 and S3). We find that the two N-terminal fragments MYC(1–88) and MYC(89–150) interact with the MYC:MAX DBD (Figure 2E) and with MYC(303–351), which immediately precedes the DBD (Figure 2B and C). In addition, the central fragments MYC(151–203), (203–255) and (256–299) also bind to the MYC:MAX DBD, while the more C-terminal MYC(303–351) does not (Fig 2E). These results were consistent regardless which one of the two components under investigation was NMR-active (Figure 2C and D). Taken together, this data clearly demonstrates binding of the MYC N-terminal fragments to the C-terminal MYC region, especially to the MYC:MAX DBD.

### Mapping of interaction sites

To map the respective binding sites of MYC fragments, we used published resonance assignments for MYC(1–88) (BMRB: 26662) and the MYC:MAX DBD (BMRB: 27571) and assigned the MYC constructs 151–255 and 256–351 ourselves (BMRB 51636 and 51635; respectively). Using standard triple resonance experiments, we assigned 100% of the backbone amide chemical shifts in both constructs (Figure S5 and S6). Assignments of MYC(151–255) and MYC(256–351) could easily be transferred to the shorter fragments MYC(151–203), MYC(203– 255), MYC(256–299) and MYC(303–351), as we found them to act as independent entities within the continuous polypeptide chain.

Binding of the MYC(1–88) and MYC(303–351) fragments could be mapped to an interaction between the conserved Mb0 (residues 15–32) and an extended stretch including the MbIV (residues 304–324) and the NLS (Nuclear Localization Signal, residues 320–328) (Figure S7). As we measured an affinity in the millimolar range, we did not further pursue this interaction. However, we cannot rule out that it is indeed biological meaningful.

Most MYC:MYC interactions identified in this study involve the structured MYC:MAX DBD (Figure 2C). Except for the basic MYC(303–351), all other fragments interact with the highly positively charged MYC:MAX DBD in our NMR binding assay (Figure 3A and Figure S3). The interaction between MYC(1–88) and the MYC:MAX DBD is reminiscent of the binding of the disordered MAX N-terminus to the DBD that we showed previously^51^. The MYC(1–88) construct is well studied with regard to MbI-dependent MYC degradation and heterotypic protein-protein interactions^15,20,27,64^. Chemical shift mapping revealed that within MYC(1–88) the Mb0 motif is the binding site for the DBD, while the MbI motif is not involved in the interaction (Figure 3B). In the MYC:MAX DBD, MYC and MAX residues within the bHLH motif show significant CSPs upon binding of the isolated Mb0 peptide, while residues in the leucine zipper are not involved in binding (Figure S8). We could further support these results by paramagnetic relaxation enhancement (PRE) data. To this end, we produced recombinant MYC(11–34) (Mb0 peptide), coupled it with the TEMPO spin label on residue C25 and recorded spectra of the MYC:MAX DBD in presence of TEMPO-labeled Mb0 in the paramagnetic (oxidized) and diamagnetic (reduced) form. The free radical of the oxidized spin label results in enhanced relaxation of nearby residues, i.e. within the binding site, and consequently in reduced peak intensities. Decreased I_ox_/I_red_ intensity ratios over a stretch of residues identifies the binding site. Our analysis showed that MYC and MAX residues especially between the basic region and helix 1 and between the loop and helix 2 experience PRE effects (Figure S8), corroborating our CSP data.

**Figure 3.**
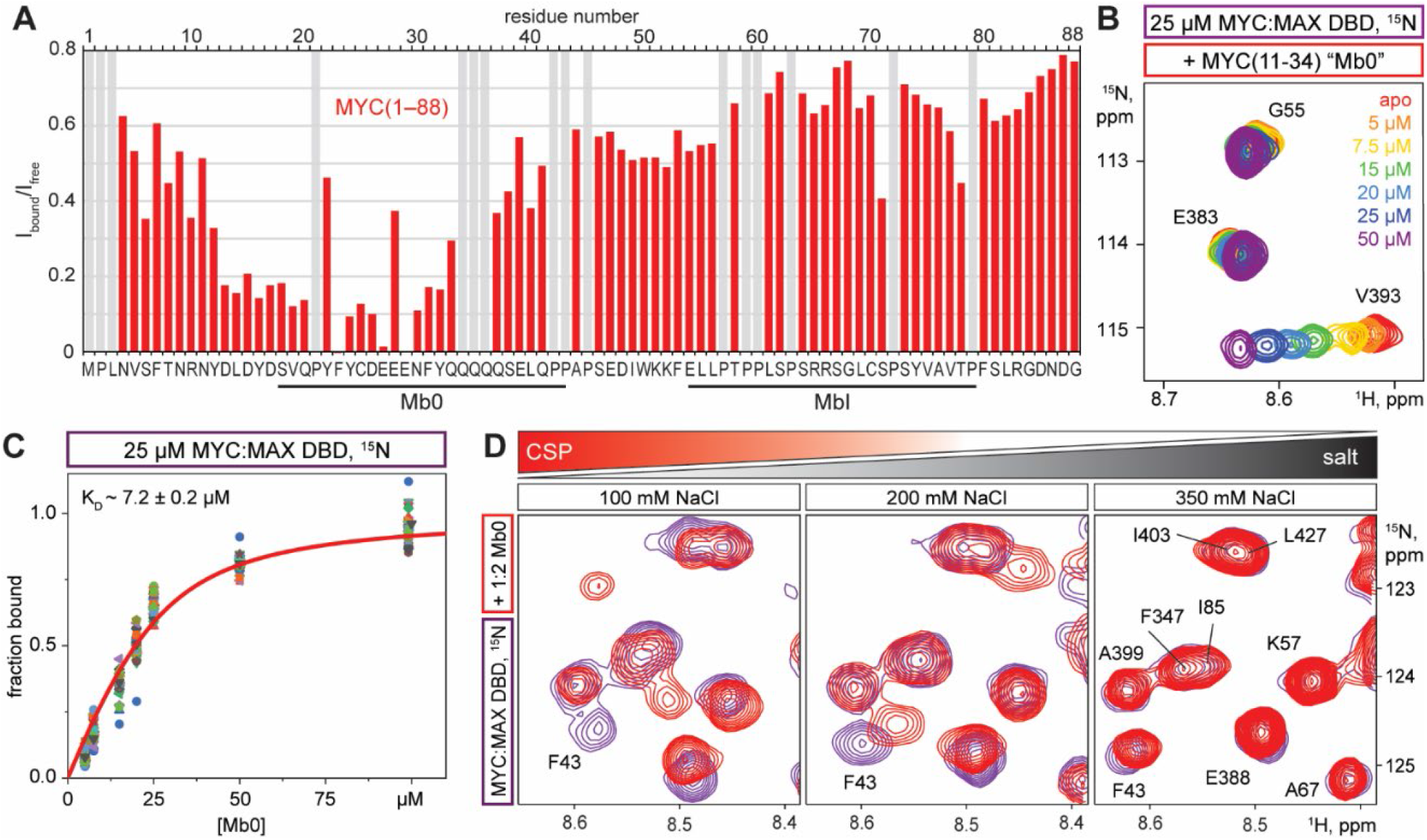
Characterization of the interaction between Mb0 and the MYC:MAX DBD. (A) Binding of MYC:MAX DBD to ^15^N-labeled MYC(1–88). Plots of I_bound_/I_free_ intensity ratios against the amino acid sequence reveal binding sites when the ratios deviate substantially from 1, as observed for residues in the conserved Mb0 motif. The MbI is not affected. (B) Zoom on ^1^H-^15^N TROSY spectra of 25 μM MYC:MAX DBD in presence of increasing concentrations of unlabeled Mb0 peptide indicated by gradual coloring from red (0 μM) to purple (50 μM). Full spectra are shown in Figure S9A. (C) Analysis of NMR titration data for the interaction shown in (B). The K_D_ ± SD is obtained from fitting the titration data globally (red line) with equation (2). (D) Zoom on ^1^H-^15^N TROSY spectra of 25 μM ^15^N-labeled MYC:MAX DBD in absence (purple) and presence of 50 μM Mb0 peptide (red) in buffer containing 100, 200 or 350 mM NaCl (from left to right).

We next performed titration experiments on NMR-active MYC:MAX DBD with unlabeled Mb0 peptide (Figure 3C and Figure S9), which revealed an affinity in the low micromolar range (K_D_ ∼7 μM; Figure 3D). In addition to Mb0 within MYC(1–88), we were able to map three additional interaction sites with the MYC:MAX DBD in the central MYC region using our new assignments (Figure S10). First, a region around the MbIIIa motif within MYC(151–203); second, the PEST region enriched in proline (P), glutamate (E), serine (S), and threonine (T) residues within MYC(203–256); and third, MbIIIb within MYC(256–299). These fragments exhibit affinities between 50 and 500 μM (Figure S11; Table 1). As we lack resonance assignments for MYC(89– 150), we were not able to identify the residues involved in binding to the MYC:MAX DBD for this MYC fragment.

**Table 1.** Summary of NMR-derived K_D_ values for MYC:MYC interactions described in this study.

| NMR-active | NMR-inactive | $K_D \pm \text{SD}$ |
| --- | --- | --- |
| MYC:MAX DBD | MYC(11–34) = Mb0 | $7.2 \pm 0.2 \mu\text{M}$ |
| MYC:MAX DBD | MYC(203–255), contains PEST | $49 \pm 1 \mu\text{M}$ |
| MYC(256–299), contains MbIIIb | MYC:MAX DBD | $361 \pm 13 \mu\text{M}$ |
| MYC(151–203), contains MbIIIa | MYC:MAX DBD | $518 \pm 16 \mu\text{M}$ |
| MYC(303–351), contains MbIV | MYC(1–88), contains Mb0+MbI | $> 1000 \mu\text{M}$<br>($1205 \pm 62 \mu\text{M}$) |

### Interactions are driven by electrostatic effects

The identified DBD-binding MYC fragments have isoelectric points (pI) below 6.1 (Figure 2B) and are, consequently, negatively charged in our NMR experiments (pH 7) and under physiological conditions. The acidic character of the MYC box motifs implies that their interactions with the MYC:MAX DBD are mainly driven by electrostatic attraction. We showed this exemplarily for binding of the DBD to Mb0 and MYC(256–299), for which we found that CSPs are reduced with increasing salt concentrations and almost completely abolished above 300 mM NaCl (Figure 3D and Figure S12A). Notably, higher salt concentrations are required to fully abrogate binding of Mb0 to the DBD, which coincides with the stronger affinity of this interaction compared to binding of MYC(256–299) to the DBD. These results support electrostatic interactions as a general mechanism for the interaction of acidic MYC fragments with the basic DBD. Indeed, the MYC and MAX bHLH-LZ motifs have high pI values of 10.1 and 9.9, respectively, whereas the interacting Mb0, MbIIIa and MbIIIb motifs have low, but comparable pI values of 3.3, 3.4 and 4.3. Nevertheless, we measure binding affinities that differ by almost two orders of magnitude, ranging from single-digit to several hundred micromolar for the Mb0 and the MbIIIb-containing MYC fragment, respectively (Figure 3D and Figure S11). Hence, other, more specific interactions based on amino acid composition or (transient) structural elements might as well contribute to binding. These considerations are supported by the observation that increasing concentrations of the polyanionic Mb0 or chloride ions (salt) do not result in similar effects on the MYC:MAX DBD. Although the MYC:MAX DBD senses increasing concentrations of salt, the extent, direction, and number of CSPs is completely different compared to the titration with Mb0 peptide (Figure S12B). Hence, the Mb0:DBD interaction that we characterized is specific and unrelated to the mere increase of negatively charges. We conclude that the observed CSP pattern on the MYC:MAX DBD is reporting on *bona fide* binding and is not a result of the gradual formation of tertiary or quaternary structure that could be attributed to the binding of chloride ions^31,65^.

### Intramolecular backfolding and DNA binding are mutually exclusive

Previously, we showed that the disordered, acidic MAX N-terminus binds to the MYC:MAX DBD in a DNA-competitive manner^51^. We thus anticipated that binding of the acidic MYC fragments to the DBD also directly competes with DNA binding. To this end, we performed competition experiments by NMR with MYC:MAX DBD, DNA and recombinant Mb0 peptide (MYC(11–34)) or MYC(256–299). For the former, we coupled an NMR-active ^13^C-labeled methyl group to a reactive cysteine (C25) via a disulfide linkage. The resulting S-^13^C-methylthiocysteine (^13^C-MTC) has been shown to possess favorable NMR properties for structural and functional studies, even for large proteins and complexes^66^. Both Mb0(^13^C-MTC-25) and ^15^N-labeled MYC(256–299) do not bind isolated E-Box DNA, but they clearly interact with the MYC:MAX DBD, resulting in CSPs between the free and the bound state (Figure 4A, Figure S13A). However, in presence of MYC:MAX DBD and E-Box DNA, resonances of the peptides appear close to or at the same chemical shift as for the free peptides, indicating displacement of the peptides from DBD by the DNA. For MYC(259–299), this was achieved by equimolar addition of DNA (Figure S13A), while for the Mb0 peptide a 5-fold excess of DNA was required to obtain full displacement (Figure 4A). This is in line with the higher affinity of the Mb0 for the DBD as that of MYC(256– 299) (Table 1). These results argue that in presence of saturating amounts of E-Box DNA, the MYC:MAX DBD can no longer interact in *trans* with the MYC boxes, likely because binding sites in the DBD are occupied by DNA, and, consequently, the MYC fragments appear as free proteins. Based on these experiments, which assessed the DNA-competitive interaction of MYC box motifs and the DBD in *trans*, i.e. as an intermolecular interaction, we aimed for an experimental setup to confirm the observed effect in an intramolecular setting. To this end, we designed MYC:MAX complexes, where we artifically fused the Mb0, MbIIIa or MbIIIb peptides directly to the MYC bHLH-LZ motif (Figure 4B and Figure S13B). A short linker between the motifs ensures sufficient mobility of the peptides to engage in binding to the DBD. We then compared these constructs with MYC(352–437):MAX(2–160), that lacks any DBD-interacting MYC box, in our SPR assay using E-Box and mutant E-Box 2 DNA (Figure 4B-D, Figure S14). We found that the Mb0 has the largest effects, both on reducing overall affinity for DNA and on providing selectivity for the consensus E-Box (Figure 4C and D). The effects for the MbIIIa- and MbIIIb-linked MYC:MAX DBDs were less pronounced, which coincides with the lower affinities that we measured for binding of the MbIIIa- and MbIIIb-containing MYC fragments to the MYC:MAX DBD compared to the high affinity for binding of Mb0 to the DBD (Table 1).

**Figure 4.**
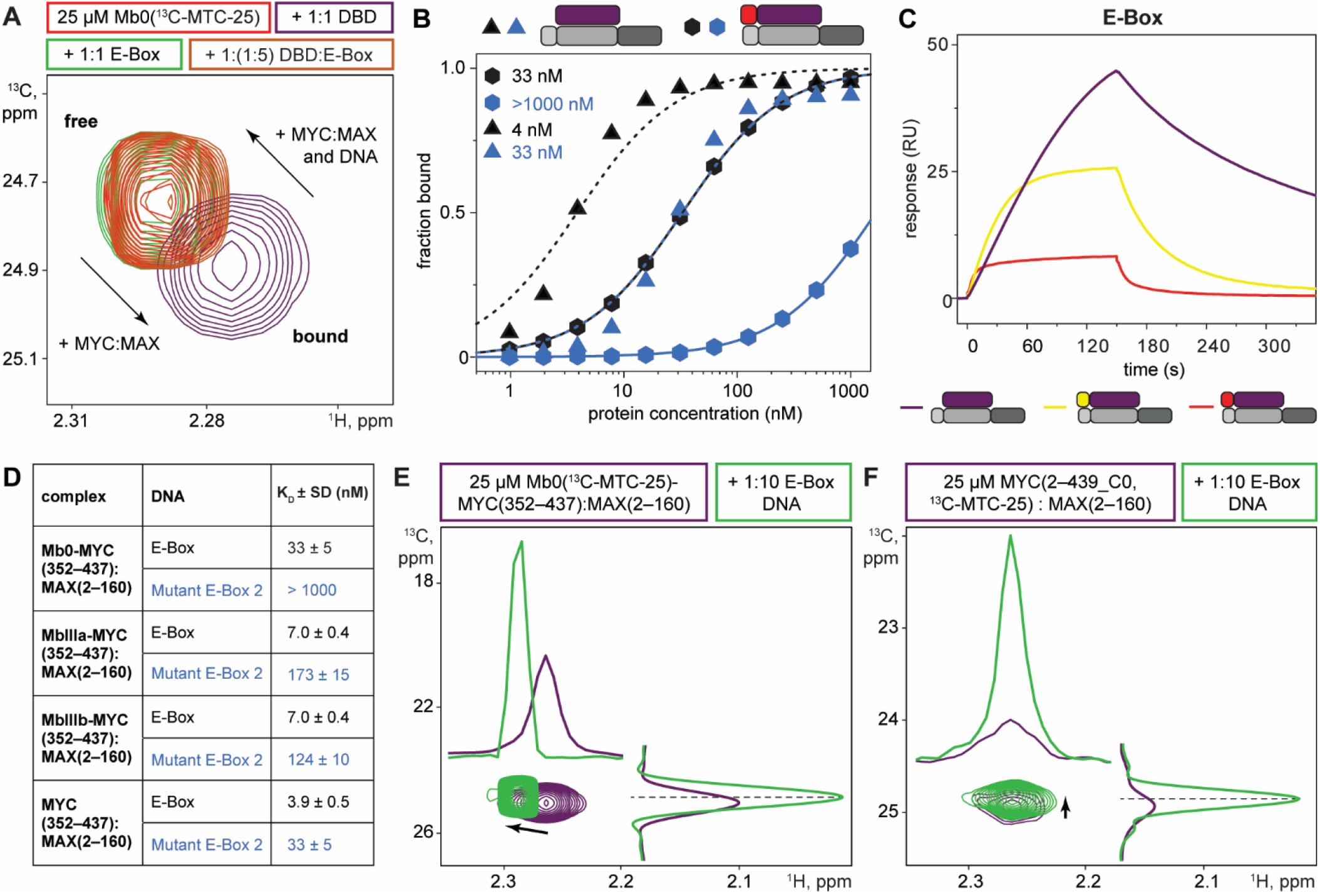
MYC boxes and E-Box DNA compete for the same binding site on the MYC:MAX DBD. (A) ^1^H-^13^C HMQC spectra of 25 μM Mb0, that is ^13^C-Methylthiocystein (MTC)-labeled at residue 25, in its free form (red) or in presence of equimolar amounts of E-Box DNA (green) and MYC:MAX DBD (purple) or in a 1:1:5 ratio (brown) with DBD and E-Box-DNA. (B) Steady-state analysis for the binding of MYC(352–437):MAX(2–160) (triangles, dotted lines) and Mb0-MYC(352–437):MAX(2–160) (hexagons, solid lines) to immobilized E-Box (black) and mutant E-Box 2 DNA (blue). K_D_ ± SD values are obtained from a global fit with equation (3), assuming a 1:1 binding model. (C) Representative SPR sensorgrams for binding of MYC(352– 437):MAX(2–160) (purple), as well as of Mb0-(red) or MbIIIa-(yellow) fusions thereof, to E-Box DNA at 3.9 nM protein concentration, illustrating the accelerating effect of the directly tethered MYC boxes on the dissociation rate. (D) Summary of K_D_ values ± SD from equilibriums fits in (B) and Figure S13B. Fold-change values are with respect to MYC(352–437):MAX(2–160). (E, F) ^1^H-^13^C HMQC spectra of (E) 25 μM Mb0-MYC(352–437):MAX(2–160) and (F) 25 μM full-length MYC:MAX that are ^13^C-MTC-labeled at MYC residue 25 in absence (purple) and presence (green) of a 10-fold molar excess of E-Box DNA. One-dimensional cross-sections in the ^1^H and ^13^C dimension are shown to illustrate the increase in peak intensity upon addition of DNA.

### Conquer full-length MYC:MAX by NMR

Finally, we attempted to demonstrate by NMR intramolecular backfolding of MYC box motifs to the DBD in the native, full-length proteins. The long unstructured regions in MYC, but also in MAX (Figure S1), pose a severe difficulty for structural studies. In addition, the size of the full-length MYC:MAX complex of about 66.5 kDa and the combination of both disordered regions and folded parts make it challenging for NMR. In initial attempts to make full-length MYC:MAX amenable for NMR, we refolded uniformly ^13^C, ^15^N-labeled MAX with unlabeled MYC. However, we only observed resonances that correspond to the disordered MAX C-terminus, while resonances for the MAX bHLH-LZ were not detectable. This is likely due to line broadening stemming from unfavorable relaxation mechanisms in the full-length protein complex.

We thus turned to post-translational isotope labeling and employed the ^13^C-MTC appproach used earlier for the Mb0. We designed a “Cys-light” MYC construct, in which all cysteine residues except C25 were mutated to alanine (MYC(2–439_C0, C25)), reconstituted dimers with full-length MAX, and incubated the complexes with ^13^C-MMTS (^13^C-methyl methanethio-sulfonate). As MAX is devoid of any cysteine, we achieved 100% labeling on the single remaining Cys25 (^13^C-MTC-25) in the Cys-light MYC without any side reaction.

In addition, we similarly labeled the Mb0-MYC(352–437):MAX(2–160) construct. As expected, methyl-TROSY spectra of the two ^13^C-MTC-labeled MYC:MAX complexes display a single peak (Figure 4E and F), that showed CSPs upon addition of E-Box DNA. Strikingly, the peak shifts are accompanied by a strong increase in peak intensity (compare 1D cross-sections through the peaks in both ^1^H and ^13^C). We attribute this observation to a order-to-disorder transition of the ^13^C-MTC-labeled Mb0 from a relatively rigid DBD-bound form in the absence of DNA to a more flexible unbound form in presence of DNA, i.e. when DNA displaces Mb0 from the DBD binding site. In the latter case, the Mb0 behaves more like a free, small disordered peptide resulting in faster tumbling and increased peak intensity, although it is still tethered to the rest of the MYC protein in the large MYC:MAX complex.

Additionally, we could displace DBD-bound Mb0 in a concentration dependent manner with excess of MAX N-terminal peptide. The pS2, pS11 double-phosphorylation on the MAX N-terminus has recently been shown to increase its intrinsic affinity for the MYC:MAX DBD^51^. We thus assumed competition of the MAX N-terminus and the Mb0 for the same DBD binding site and performed the NMR competition experiment with phosphorylated MAX(2–21). Indeed, we observed concentration-dependent CSPs and peak intensity increases for the ^13^C-MTC-labeled Mb0-MYC:MAX complex upon addition of phosphorylated MAX(2–21) (Figure 13C). Expectedly, competition can also be achieved with excess of free Mb0 peptide. For these experiments we used a Mb0 peptide, whose reactive Cys25 thiol was blocked by alkylation with iodoacetamide to prohibit disulfide shuffling between the free and the DBD-linked, ^13^C-MTC-labeled Mb0 (Figure 13D). We thus conclude that E-Box DNA as well as high-affinity DBD-interacting motifs in MYC and MAX efficiently compete for the same binding site on the DBD, also in context of native, full-length MYC:MAX.

All in all, we could showcase that the transcription factor complex MYC:MAX, which is regarded challenging due to it large size and mainly disordered nature, is amenable to NMR studies. Our NMR results on full-length MYC:MAX corroborate our mechanistic SPR studies and NMR experiments on isolated MYC fragments. Together, this data suggests intramolecular backfolding of the N-terminal and central MYC box motifs onto the C-terminal DBD, although they are separated by up to 300 amino acids in the native MYC protein. This backfolding mechanism is probably relevant for regulating DNA binding as well as heterotypic protein-protein interactions in cellular settings, thereby providing an additional layer of transcriptional regulation.

## Discussion

### Electrostatic interactions drive IDR:DBD binding in the MYC:MAX complex

In this study, we identified and mapped interactions within the transcription factor MYC, one of the key human oncoproteins. Thereby, we not only observe interactions between intrinsically disordered regions (IDRs) but also between IDRs and the folded DNA binding domain (DBD) that is formed by heterodimerization of the MYC and MAX bHLH-LZ motifs. In a cellular setting, these interactions can occur intramolecularly in a single MYC protein (in *cis*) or intermolecularly between two MYC molecules (in *trans*). We will discuss first the intermolecular aspect before turning to the consequences of intramolecular binding.

Closer examination of the interacting motifs indicates that the observed binding events are mainly driven by electrostatic attraction between the positively charged (basic) DBD and negatively charged (acidic) IDRs. Our findings for MYC are reminescent of other transcription factors, for which intramolecular interactions between IDRs and DBDs have been reported in the past. These include the p53 family^67^, bZIP (CCAAT/ Enhancer-binding proteins, C/EBPs)^68^, bHLH-LZ TFs^51^ and Forkhead box (FOX) TFs^69,70^. Notably, their IDRs have low isoelectric points (pI < 4), which renders these regions negatively charged under physiological conditions and thus enables interactions with their respective positively charged DBDs (pI > 9). The results that we present in this study for MYC support electrostatics-driven interactions between acidic disordered regions and basic DBDs as an important and common regulatory mechanism that is shared among different TF families. Interestingly, the MYC:MAX DBD also interatcs with folded, negatively charged domains as was shown for the interaction with the INI1/hSNF5 RPT1 (pI = 4.1)^71^. Thereby, the pattern of affected MYC and MAX residues and the binding affinity (K_D_ = 44 μM) are very similar to the interaction of the various disordered Mb’s to the DBD (see below). A DNA-competitive engagement of the MYC:MAX DBD by acidic binding motifs thus seems to be a conserved mechanism for MYC interaction partners.

Albeit electrostatic interactions are important for interactions in MYC, we notice pronounced differences in the affinities of these binding events. In essence, we measured affinities in *trans* that cover a range of about three orders of magnitude (Table 1). Thereby, the weakest interaction is between the two disordered fragments MYC(1–88) and MYC(303–351) with a K_D_ in the low millimolar range. The affinities for the MbIIIa- and MbIIIb-containing fragments MYC(151–203) and MYC(256–299) to the MYC:MAX DBD are in the high micromolar range, while we find the PEST sequence and the Mb0 to interact with the DBD with double- and single-digit micromolar affinity, respectively (Table 1). Interestingly, the nuclear concentration of MYC was reported to be up to several micromolar, depending on cell stimulus and without consideration of sub-nuclear compartmentalization^72^. Consequently, the strong affinities that we measured are in the range of physiological MYC concentrations. Moreover, the design of our NMR binding experiments was such that the interactions were in *trans*, while in native full-length MYC these would be intramolecular interactions. We and others have shown that weak intermolecular binding events become significantly more affin and efficient when they are translated into an intramolecular setting^51,73,74^. We conclude that the interactions found in this study, especially between disordered MYC fragments and the folded DBD, are biologically meaningful. Further studies are necessary to dissect the influence of the Mb:DBD interactions on, for example, chromatin engagement^75^ and target gene activation on the cellular level^76^.

The interplay of weak but multivalent intermolecular interactions between IDRs or between IDRs and folded domains, as we described for MYC in this study, is reminescent of processes that manifest as liquid-liquid phase separation (LLPS) *in vitro* and *in vivo*^77,78^. Members of the bHLH-LZ family, among them MYC as a prominent example, contain regions with the predicted potential to drive phase separation^23^. Indeed, c-MYC has been shown to undergo homotypic LLPS and molecular determinants for this process have been dissected in more detail^79,80^. Additionally, LLPS-mediated target gene activtion has been demonstrated for the TFs OCT4 and GCN4^79^, TAZ/TEAD^81^, engineered OptoDroplet TFs^82^, and N-MYC^83^. Whether the herein identified MYC:MYC interactions contribute to transcriptionally active foci also for c-MYC has to be shown in future studies.

### Kinetic parameters establish a mechanistic model of DNA binding by the MYC:MAX TF

We used SPR as a first step to shed light on the biological relevance of the identified MYC:MYC interactions, as it allowed us to screen and compare MYC proteins of variable length ranging from the bHLH-LZ to the full-length protein. Since the quantitative assessment of kinetic DNA binding parameters (k_on_ and k_off_) was not possible for all the different MYC:MAX complexes used in this work, we restrict our discussion to a qualitative comparison. Importantly, we find that the disordered regions in MYC reduce the overall affinity to DNA by about 15-fold (Figure 1). A visual inspection of the SPR sensorgrams revealed that the affinity differences observed between the MYC:MAX DBD and full-length MYC:MAX are due to considerably slower association rates for the latter (Figure 1C). By contrast, the dissociation rates are comparable for both complexes. In this regard, the N-terminal MYC IDR with its more complex amino acid composition clearly does not behave like a classical D/E repeat^84^. It was recently shown that auto-inhibited D/E repeat containing proteins can have accelerated DNA association rates that facilitate target search, while their counterparts lacking the acidic repeats bind DNA more slowly^85^. In contrast, in this work we have shown that the auto-inhibited full-length MYC:MAX complex associates more slowly than the non-inhibited MYC:MAX DBD.

The MYC:MAX complex is dynamic and undergoes dissociation at low concentrations. This could potentially affect our SPR measurements, especially the data points that were recorded at very low concentrations, where a dissociation of MYC:MAX might influence binding kinetics. Dissociation constants for the MYC:MAX DBD have been determined to be in the range of about 30–200 nM^36,59,86,87^. The heterodimeric bHLH-LZ is destabilized mainly by charge repulsion of the basic regions (bRs) in MYC and MAX. Thus, charge compensation by deletion of the bRs^71^, addition of polyglutamate in *trans*^86^ or substitution of the MAX bR with an polyanionic stretch^87^ leads to significantly more stable dimers. In our work, we use full-length MAX, that harbors the acid N-terminus (residues 1–21) and the MYC(151–439) and MYC(2–439) proteins possess acidic IDRs in their N-termini as well. These polyanionic regions can also shield the electrostatic repulsion between the MYC and MAX bRs. We thus assume that our MYC:MAX complexes have dissociation constants that are significantly lower than those that were measured for the isolated MYC:MAX DBD - likely because the acidic regions in the longer MYC constructs stabilize the heterodimers compared to MYC(352–437). Consequently, even at the lowest concentration used in our SPR experiments (0.5 nM), we assume that the MYC and MAX proteins are predominantly in the heterodimeric form.

Based on our NMR and SPR data, we propose the following model of DNA engagement by the full-length MYC:MAX complex: a DNA-free MYC:MAX complex, as it would appear for example after translation in the cytosol, can adopt a “locked” state, where a MYC box (e.g. Mb0) folds back to the DBD, creating a conformational situation that is reminescent of DNA binding (Figure 5A). Once in the nucleus, DNA would have to compete with the MYC box that occupies the DBD. As a consequence, this competition slows down association of the DNA to MYC:MAX, since the MYC box has to dissociate first. The MYC box, that is now released from the DBD, is separated from the binding site by a relatively long linker, making it stochastically less likely to engange in productive encounters with a DNA-bound DBD. Consequently, full-length MYC:MAX has comparable dissociation rates as MYC(352–437):MAX(2–160) – in both complexes the dissociation from DNA is driven by the proximal MAX N-terminus that has an intermolecular K_D_ of 13.2 μM for the DBD and is attached to it via a very short linker^51^.

**Figure 5.**
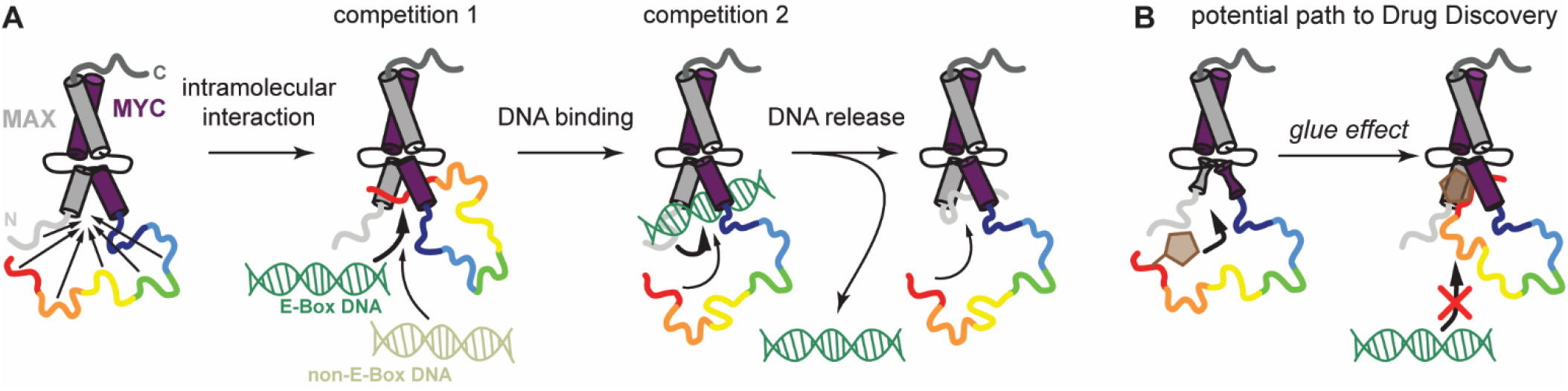
Intramolecular interactions in MYC provide DNA selectivity and offer new paths to drug discovery. (A) Schematic depiction of our current model how DBD-interacting Mb motifs influence the DNA binding behavior of MYC:MAX. See Discussion for a more detailed explanation. (B) Potential new approach to directly target MYC:MAX by exploiting the interactions identified in this work with covalent drug modalities that stabilize the MYC box : DBD interactions, resulting in further reduced DNA binding.

These considerations are supported by our SPR data for the artificial Mb-linked MYC:MAX complexes. When we tethered additional DBD-interacting motifs (i.e. Mb0, MbIIIa or MbIIIb) to MYC:MAX via short linkers (Figure 4B and Figure S13B), we observed accelerated dissociation from DNA for these complexes compared to MYC(352–437):MAX(2–160), while the association rates were similar (Figure 4C), as concluded from a qualitative evaluation of the SPR sensorgrams. We rationalize this change from dissociation rate-driven affinity reduction as observed for Mb- linked MYC:MAX to association rate-driven affinity reduction as observed for full-length MYC:MAX with (1) the longer “spacers” between the MYC box motifs and the DBD in the full-length protein and (2) a redundancy in the Mb:DBD interactions. In full-length MYC, a DBD-interacting MYC box can easily be displaced intramolecularly by another one. Hence, DNA has to compete not only with one component but many, which slows down the binding process of the DNA to the DBD.

### New avenues for MYC targeting in drug discovery

Capitalizing on the herein identified and characterized interactions could provide new starting points for the identification of novel modalities to directly target MYC in drug discovery. Low molecular weight compounds could act as molecular glues that strengthen the Mb0:DBD interaction and lock MYC in a conformation that is less efficient in DNA binding (Figure 5B). Eventually, covalent approaches can even utilize the reactive Cys25 in Mb0. Our NMR data with unmodified, ^13^C-MTC-, acetamide- and TEMPO-labeled Mb0 suggests that the DBD in principal tolerates covalent modifications on Cys25 (Figure S15). Although the mentioned modifications weaken the Mb0:DBD interaction to some extent, structure-guided medicinal chemistry efforts might be able to pick up favorable interactions, which could increase the affinity. Ultimately, these efforts would result in a “locked” MYC:MAX whose binding to non cognate DNA is further reduced, thereby lowering the oncogenic potential of MYC.

## ASSOCIATED CONTENT

### Supporting Information

The following files are available free of charge.

Additional information, analysis and results as well as a comprehensive list of oligonucleotides and protein constructs used in this study (PDF).

## ACKNOWLEDGMENT

We thank César Fernandez for assistance with NMR experiment setup and analysis and Christelle Henry for maintenance of the NMR infrastructure. We would also like to thank Felix Freuler and Simon Hänni for excellent molecular biology support.

