## Supporting_Information for "Intrinsically disordered regions in the transcription factor MYC:MAX modulate DNA binding via intramolecular interactions"

[1] Novartis - Biomedical Research, Novartis Campus , CH-4056 Basel, Switzerland.
[2] current adress: Ridgeline Discovery, CH-4057 Basel, Switzerland
[3] ETH Zürich, CH-8093 Zurich, Switzerland

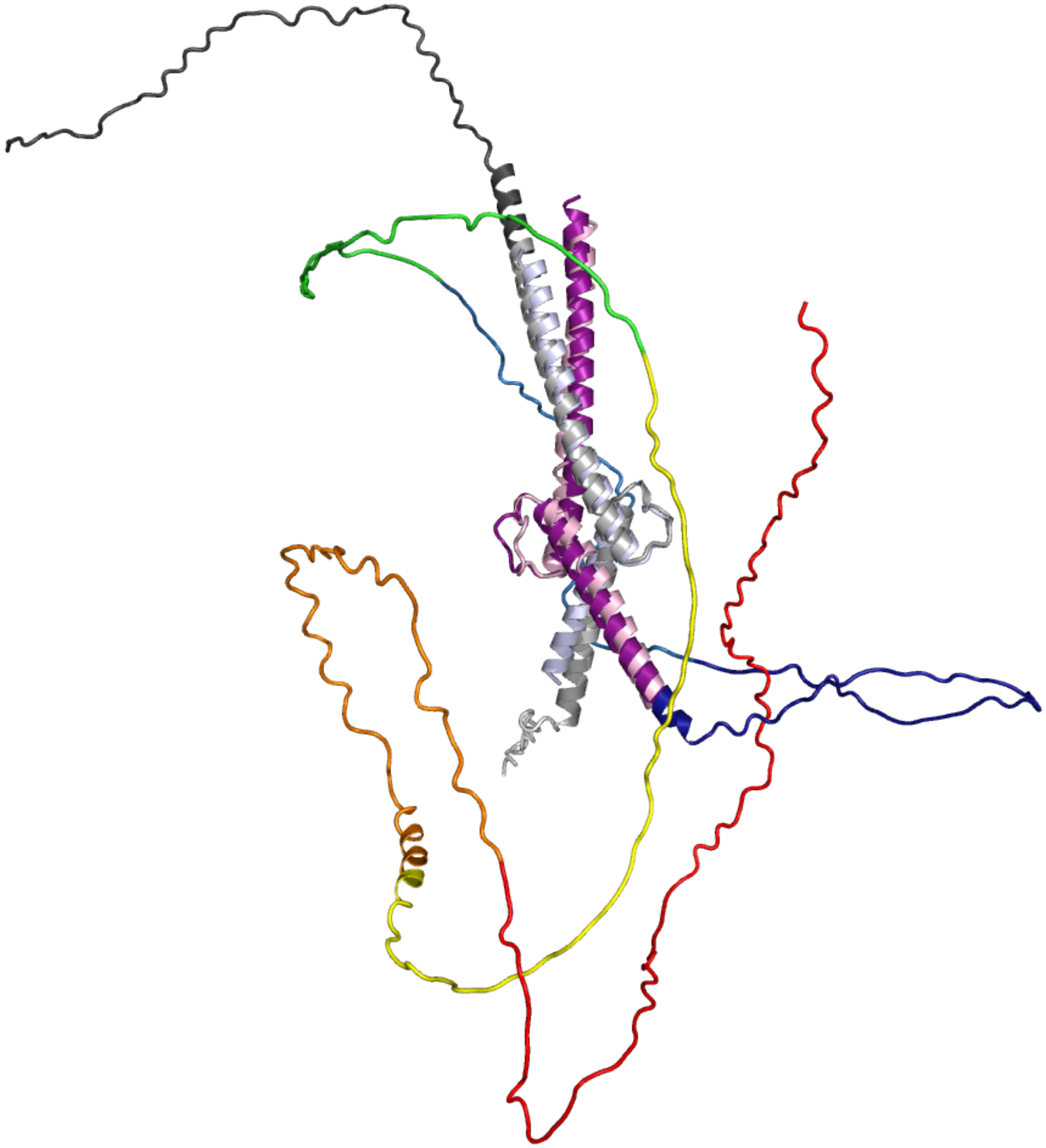

**Figure S1. MYC and MAX proteins are mainly disordered**. Model of the DNA-free MYC:MAX heterodimer as predicted by AlphaFold2 [1]. MYC is in rainbow colors according to the fragments used in this study: 1–88 (red), 89–150 (orange), 151–203 (yellow), 203–255 (green), 256–299 (blue), 303–351 (dark blue), and 352–437 (purple). MAX is colored in different shades of gray: the N-terminus (1–21) is in light gray, the bHLH-LZ (22–102) is in gray, and the C-terminus (103–160) is in dark gray. The crystal structure of the apo MYC:MAX DBD is shown for comparison (PDB: 6G6K), with MYC in pink and MAX in light blue[2].

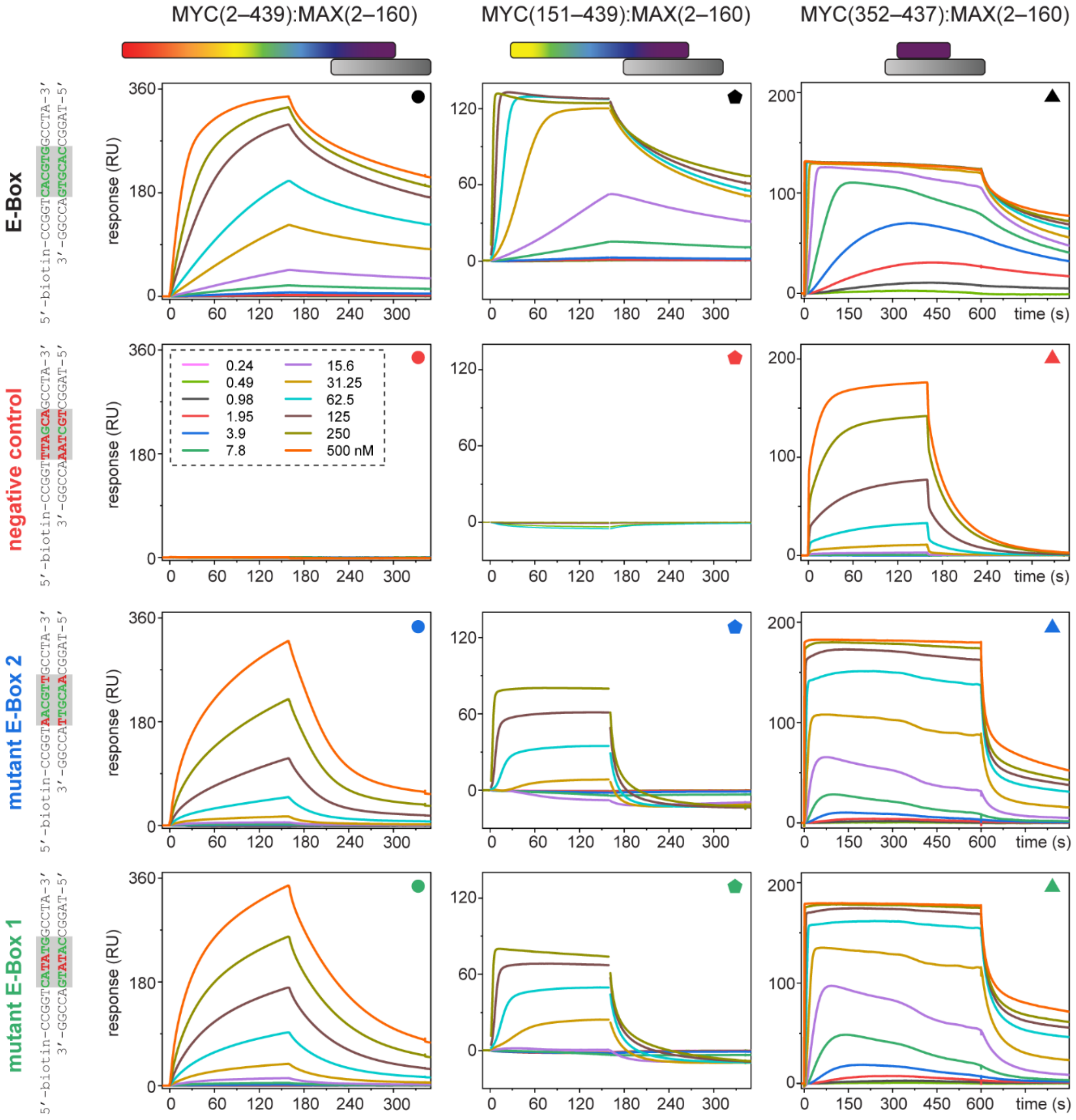


**Figure S2**. **Disordered MYC regions lead to lower DNA binding affinity and slower association.** SPR sensorgrams for the binding of full-length MYC(2–439):MAX(2–160) (left), MYC(151–439):MAX(2–160) (middle) and MYC(352–437):MAX(2–160) (right) to immobilized E-Box, negative control, and mutant E-Box DNA (from top to bottom). Altered nucleotides in the core hexameric consensus sequence are shown in red.

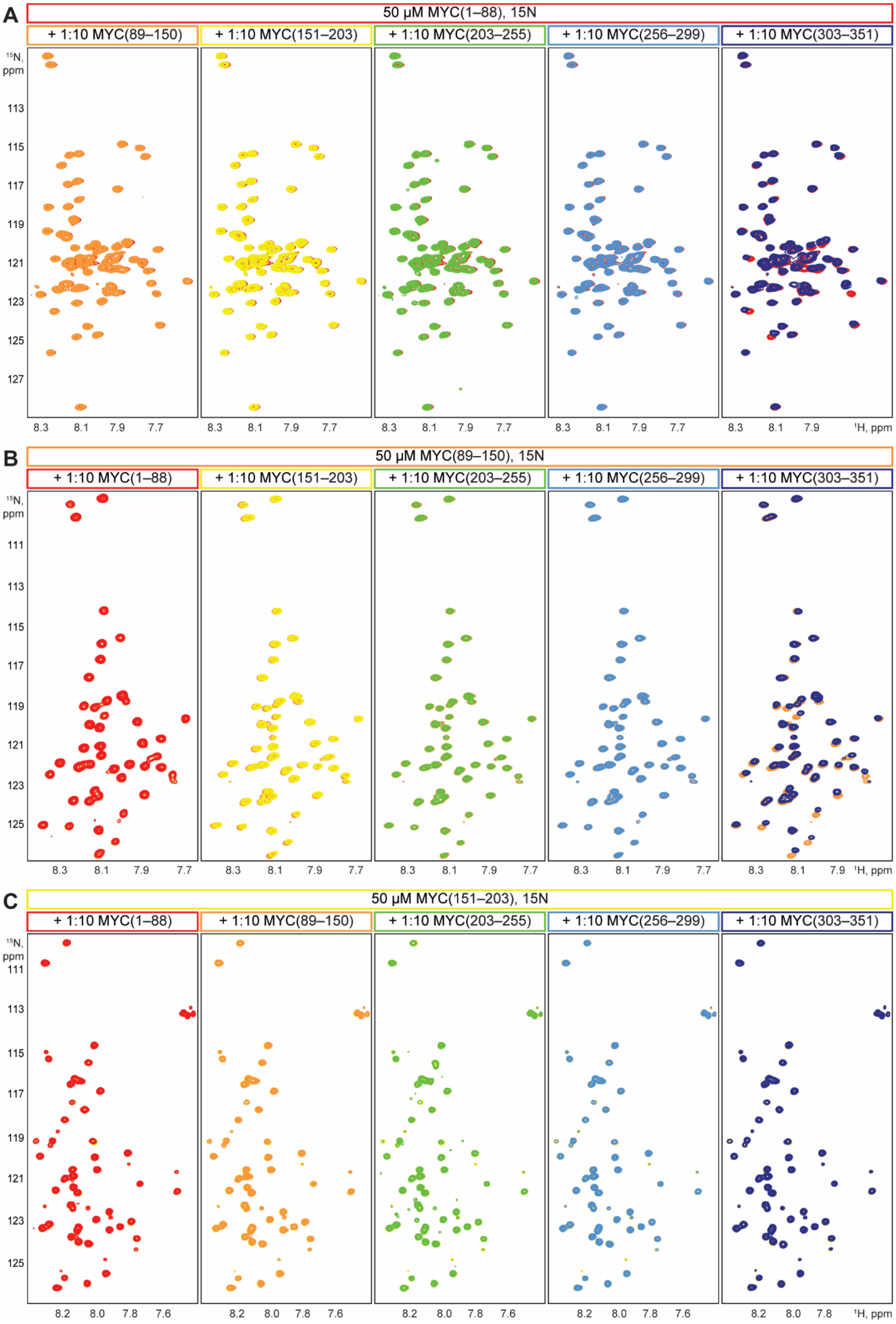
A
50 μM MYC(1–88), 15N
+ 1:10 MYC(89–150)
+ 1:10 MYC(151–203)
+ 1:10 MYC(203–255)
+ 1:10 MYC(256–299)
+ 1:10 MYC(303–351)
15N, ppm
113
115
117
119
121
123
125
127
8.3 8.1 7.9 7.7
1H, ppm
B
50 μM MYC(89–150), 15N
+ 1:10 MYC(1–88)
+ 1:10 MYC(151–203)
+ 1:10 MYC(203–255)
+ 1:10 MYC(256–299)
+ 1:10 MYC(303–351)
15N, ppm
111
113
115
117
119
121
123
125
8.3 8.1 7.9 7.7
1H, ppm
C
50 μM MYC(151–203), 15N
+ 1:10 MYC(1–88)
+ 1:10 MYC(89–150)
+ 1:10 MYC(203–255)
+ 1:10 MYC(256–299)
+ 1:10 MYC(303–351)
15N, ppm
111
113
115
117
119
121
123
125
8.2 8.0 7.8 7.6
1H, ppm

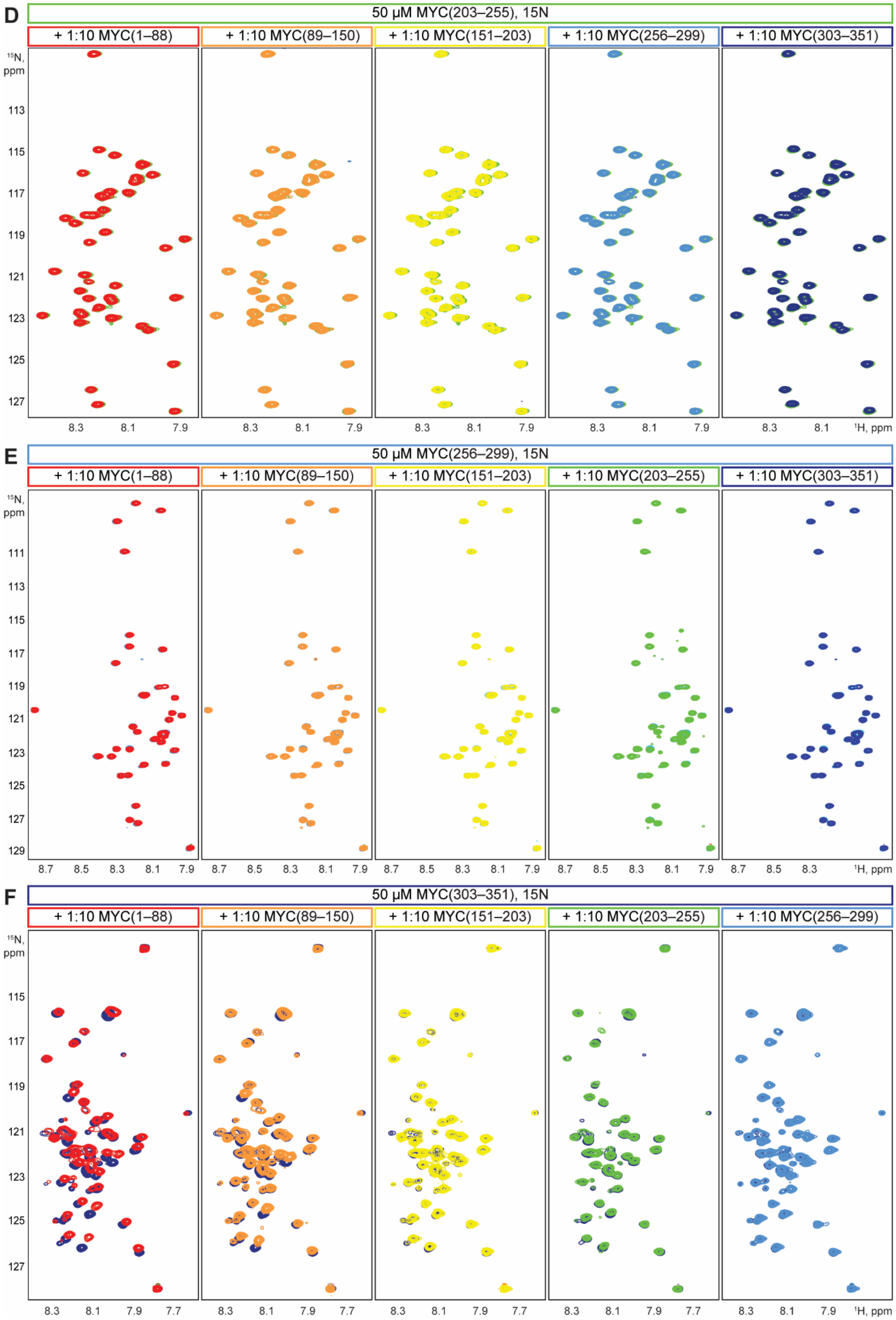
D
50 μM MYC(203–255), 15N
+ 1:10 MYC(1–88)
+ 1:10 MYC(89–150)
+ 1:10 MYC(151–203)
+ 1:10 MYC(256–299)
+ 1:10 MYC(303–351)
¹⁵N, ppm
113
115
117
119
121
123
125
127
8.3
8.1
7.9
¹H, ppm
E
50 μM MYC(256–299), 15N
+ 1:10 MYC(1–88)
+ 1:10 MYC(89–150)
+ 1:10 MYC(151–203)
+ 1:10 MYC(203–255)
+ 1:10 MYC(303–351)
111
113
115
117
119
121
123
125
127
129
8.7
8.5
8.3
8.1
7.9
F
50 μM MYC(303–351), 15N
+ 1:10 MYC(1–88)
+ 1:10 MYC(89–150)
+ 1:10 MYC(151–203)
+ 1:10 MYC(203–255)
+ 1:10 MYC(256–299)
115
117
119
121
123
125
127
8.3
8.1
7.9
7.7

**Figure S3. Interactions between disordered MYC regions:** (A) $^{1}H$-$^{15}N$ HSQC spectra of 50 μM $^{15}N$-labeled MYC(1–88) in absence (red) or presence of a 10-fold molar excess of either MYC(89–150) (orange), MYC(151–203) (yellow), MYC(203–255) (green), MYC(256–299) (blue) or MYC(303–351) (dark blue). Presence of CSPs indicates binding. The small signal dispersion in the $^{1}H$-dimension for all MYC fragments is characteristic for their intrinsic disorder. (B-F) as in (A), but for (B) $^{15}N$-labeled MYC(89–150), (C) $^{15}N$-labeled MYC(151–203), (D) $^{15}N$-labeled MYC(203–255), (E) $^{15}N$-labeled MYC(255–299), and (F) $^{15}N$-labeled MYC(303–351). Spectra in presence of a 10-fold molar excess of MYC(1–88) are in red.

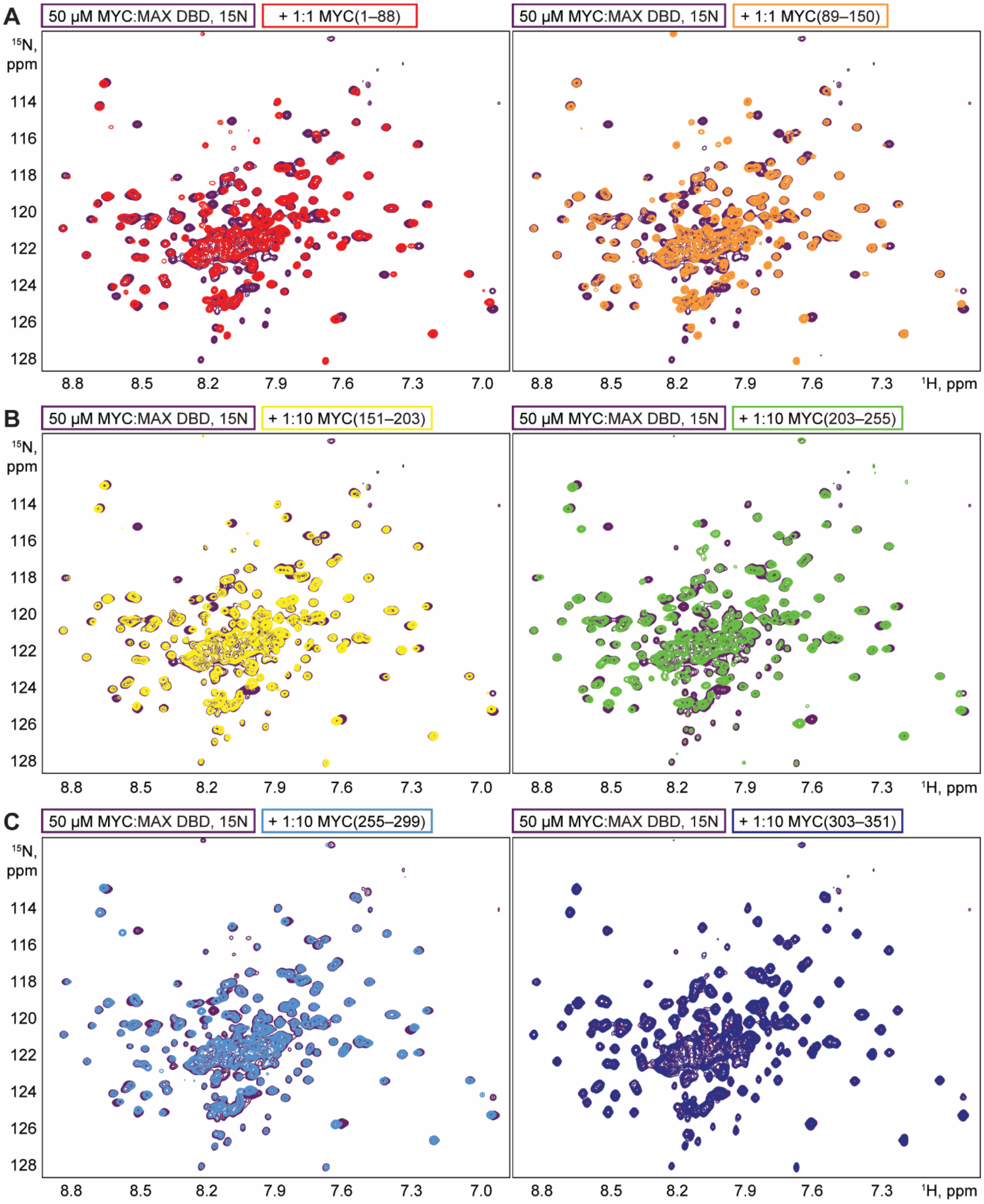


**Figure S4. The folded MYC:MAX DBD interacts with disordered MYC fragments**: (A) $^{1}$H-$^{15}$N TROSY spectra of 50 μM $^{15}$N-labeled MYC:MAX DBD, in absence (purple) or presence of an equimolar amount of either MYC(1–88) (red) or MYC(89–150) (orange). Presence of CSPs and decreases in peak intensity indicate binding. The large dispersion in the $^{1}$H dimension is characteristic for folded proteins. (B) as in (A), but with a 10-fold molar excess of either MYC(151–203) (yellow) or MYC(203–255) (green). (C) as in (A), but with a 10-fold molar excess of either MYC(256–299) (blue) or MYC(303–351) (dark blue).

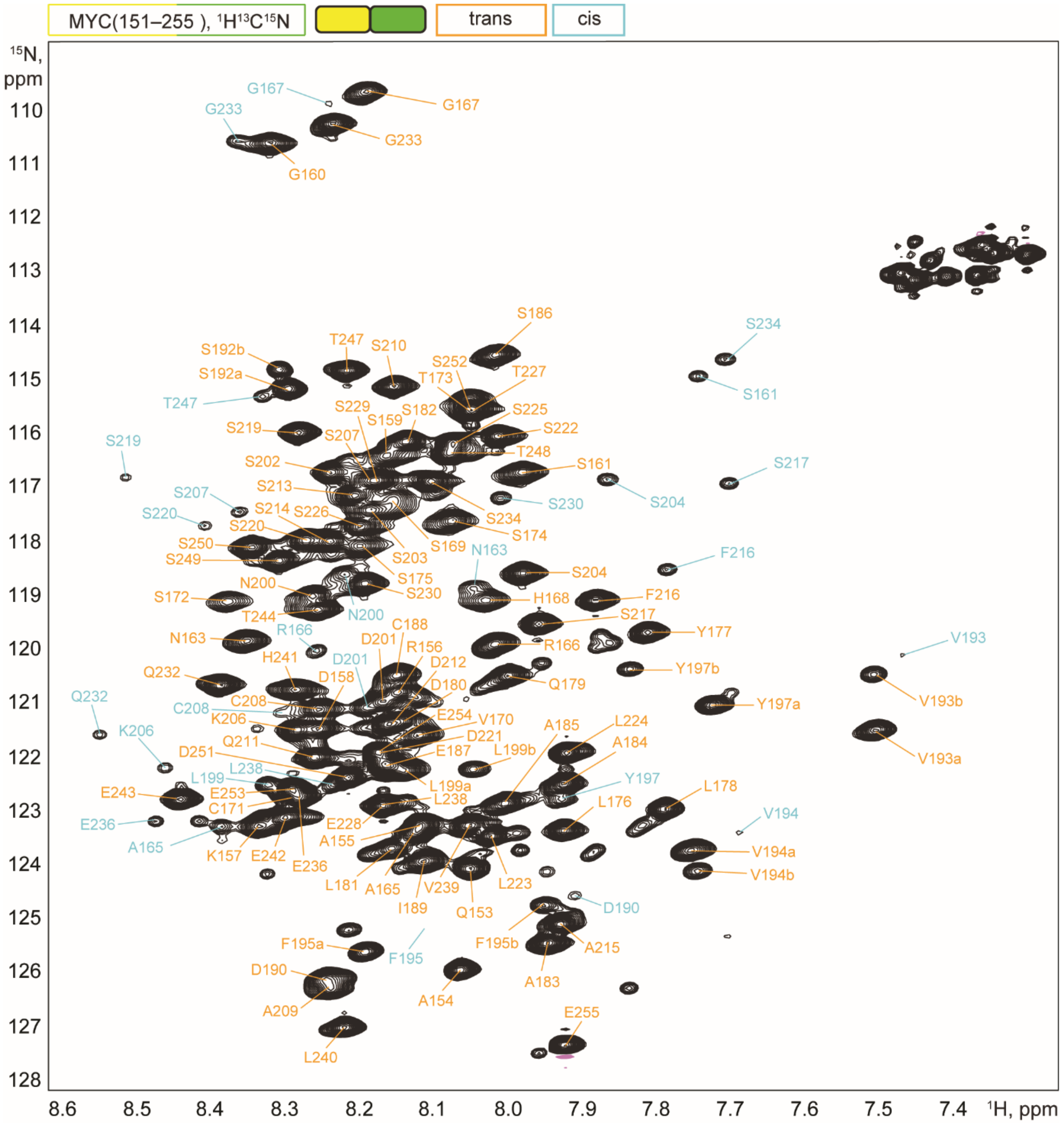


**Figure S5. Assignment of MYC(151–255):** $^{1}$H-$^{15}$N HSQC spectrum of 665 µM $^{13}$C, $^{15}$N-labeled MYC(151–203). Resonance assignments are indicated in orange for the major form with all prolines in *trans*. For various prolines a less populated *cis* conformation at the amide bond is observed, which gives rise to minor peaks for some residues. Assignments for this minor conformation are colored cyan. Assignments are deposited under the BMRB accession number 51636.

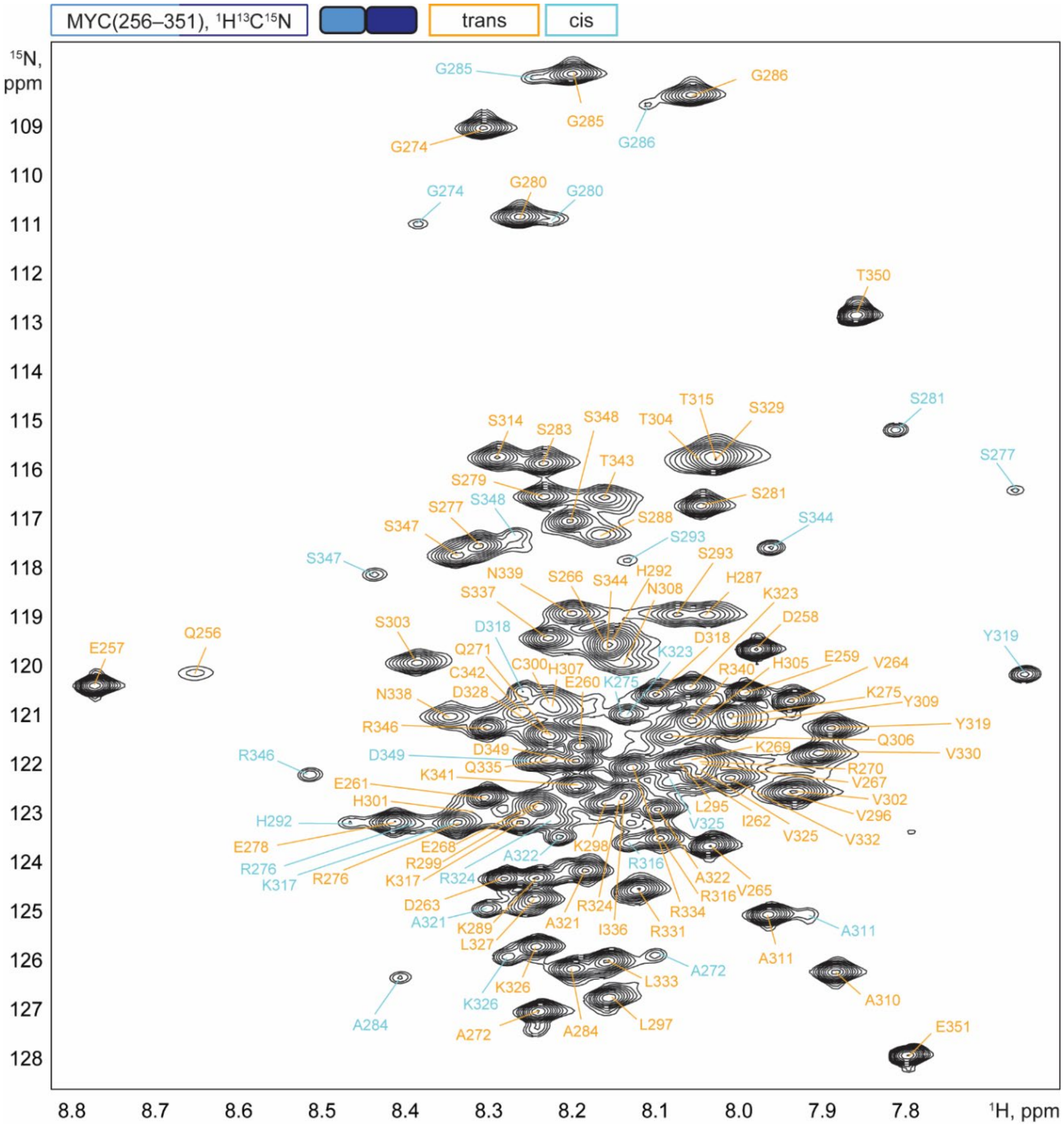


**Figure S6. Assignment of MYC(256–351):** $^{1}$H-$^{15}$N HSQC spectrum of 900 µM $^{13}$C, $^{15}$N-labeled MYC(256–351). Resonance assignments are indicated in orange for the major form with all prolines in *trans*. For various prolines a less populated *cis* conformation at the amide bond is observed, which gives rise to minor peaks for some residues. Assignments for this minor conformation are colored cyan. Assignments are deposited under the BMRB accession number 51635.

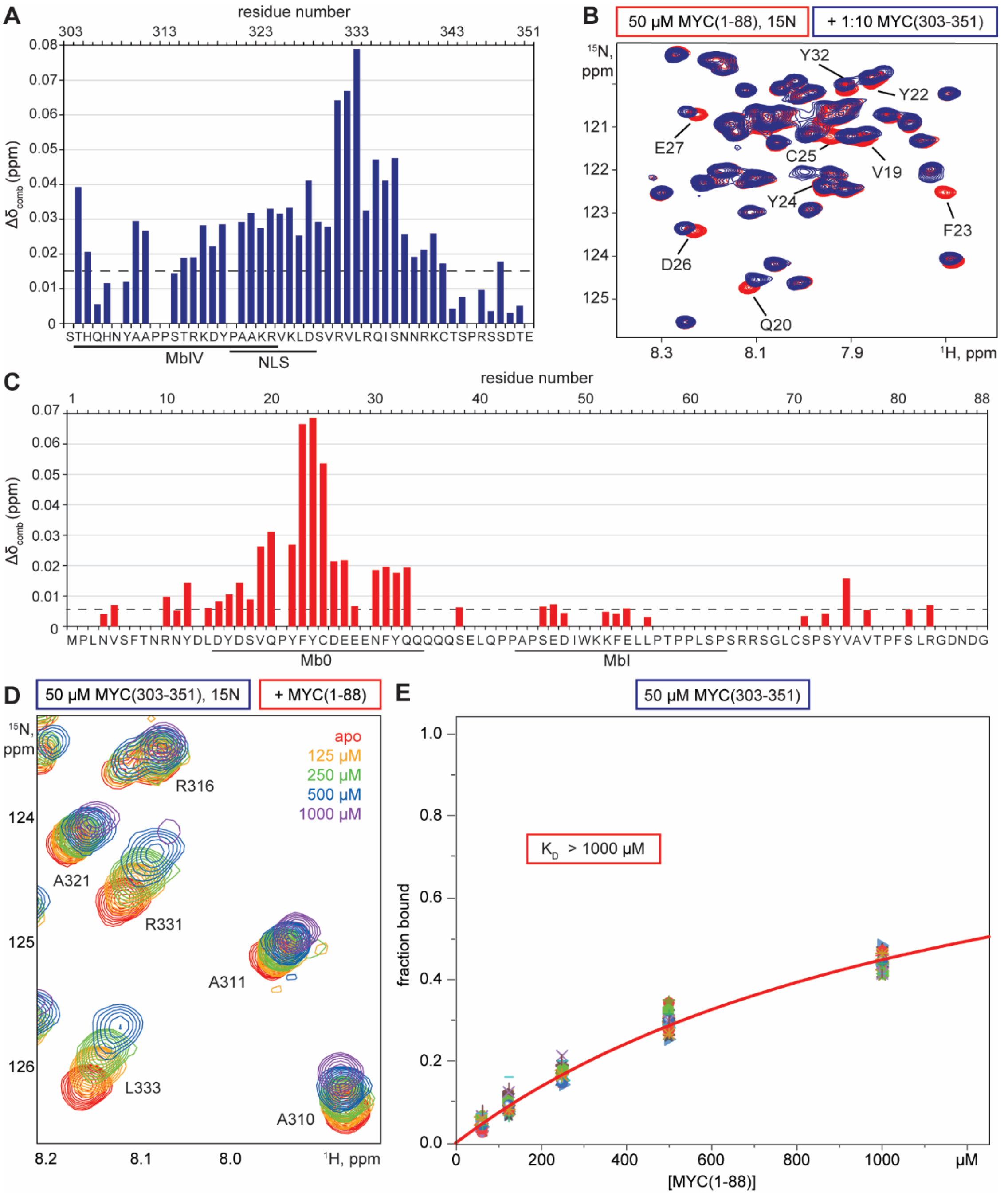


**Figure S7. The MYC fragments (1–88) and (303–351) interact weakly via conserved motifs.** (A) Quantitative analysis of CSPs observed on MYC(303–351) in presence of MYC(1–88). The binding site extends C-terminally over the conserved MbIV motif and the nuclear localization signal (NLS). (B) Zoom on $^{1}$H-$^{15}$N HSQC spectra of 50 µM $^{15}$N-labeled MYC(1–88) in absence (red) and presence (blue) of a 10-fold molar excess of MYC(303–351). Assignments are indicated for resonances showing CSPs. (C) as in (A), but for CSPs observed on MYC(1–88) in presence of MYC(303–351). Affected residues cluster in the conserved Mb0 motif, while MbI is not involved in binding. (D) Zoom on $^{1}$H-$^{15}$N HSQC spectra of 50 µM $^{15}$N-labeled MYC(303–351) with increasing concentrations of MYC(1–88) indicated by gradual coloring from red (0 µM) to purple (1000 µM). (E) Analysis of NMR titration data for the interaction in (D). The $K_D$ is obtained from a global fit (red line) with equation (2) and higher than the highest ligand concentration used ($K_D$ > 1000 µM).

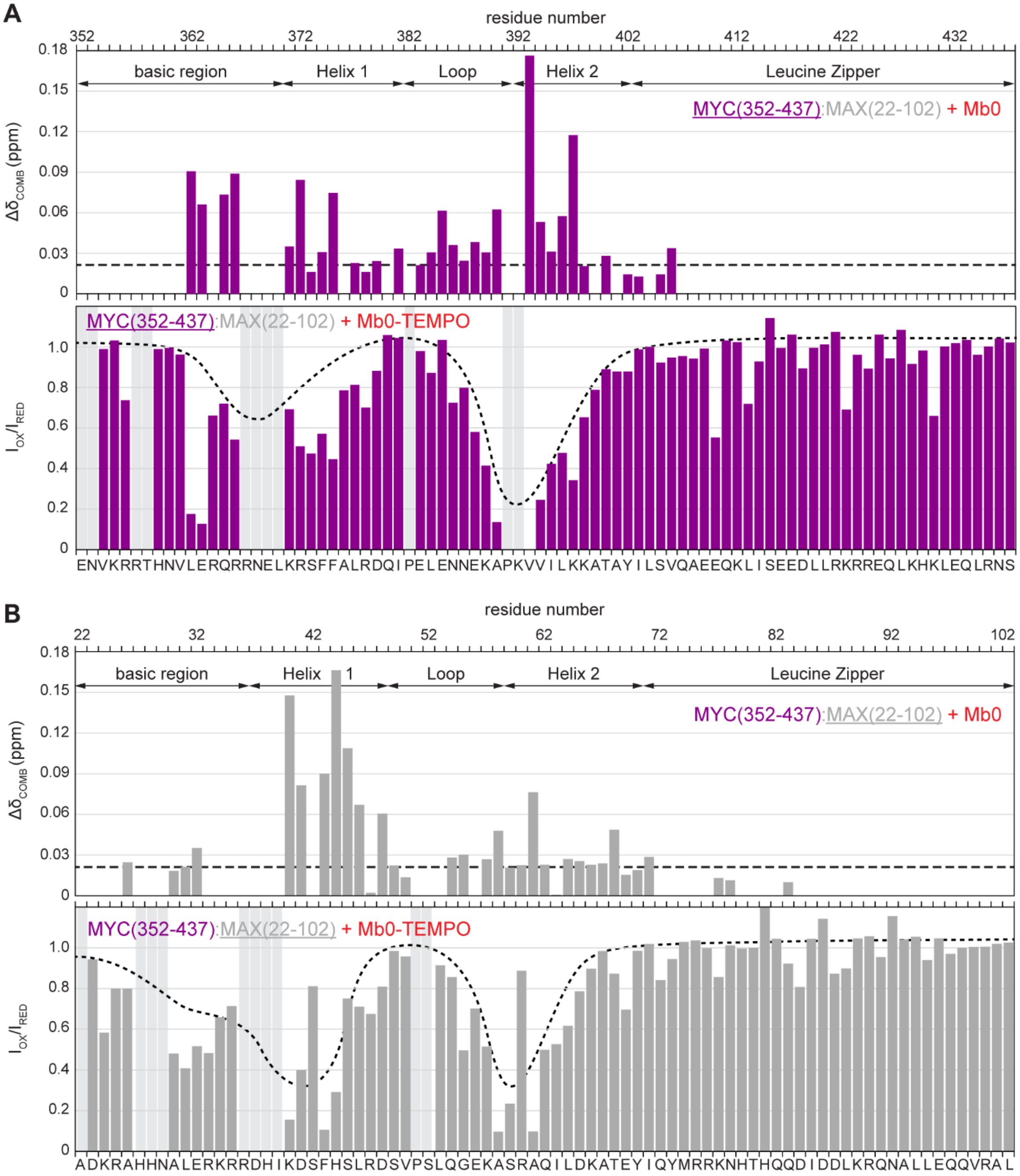


**Figure S8. Mb0 interacts with the basic region and the HLH motif within the MYC:MAX DBD.** (A, B) Quantitative analysis of CSPs (top) and PRE effects (bottom) observed for bHLH-LZ residues in MYC (A) and MAX (B) in presence of Mb0. Residues affected by binding localize to the basic region and the HLH motif. The leucine zipper is not involved in binding to Mb0. The dashed lines indicate above which value CSPs are significant (see Materials and Methods). The dotted lines illustrate the reduction in signal intensity due to PRE effects upon binding of spin-labeled Mb0 (Mb0-TEMPO) to the DBD. Light gray bars indicate missing assignments.

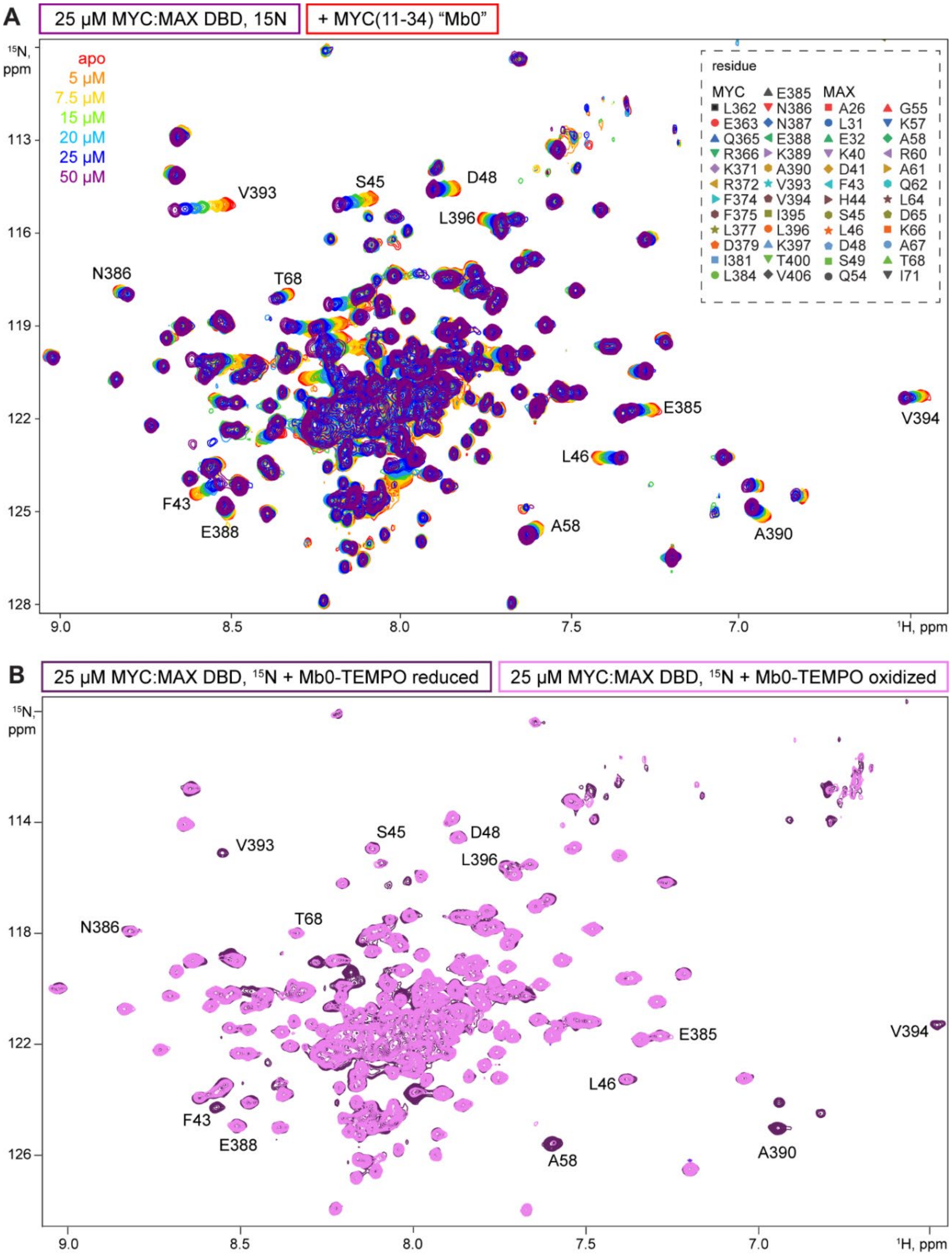


**Figure S9. Mb0 binds the MYC:MAX DBD with low micromolar affinity**. (A) $^{1}H$-$^{15}N$ TROSY spectra of 25 µM $^{15}N$-labeled MYC:MAX DBD with increasing concentrations of MYC(11–34), Mb0, indicated by gradual coloring from red (0 µM) to purple (50 µM). Assignments for some resonances are indicated. The inset lists residues whose resonances were used for $K_D$ determination. (B) $^{1}H$-$^{15}N$ TROSY spectra of 25 µM $^{15}N$-labeled MYC:MAX DBD with equimolar amounts of TEMPO-labeled Mb0 in the reduced (purple) and oxidized (pink) form. The spin label induces PRE effects on resonances in the bHLH motif, as shown in FigureS8.

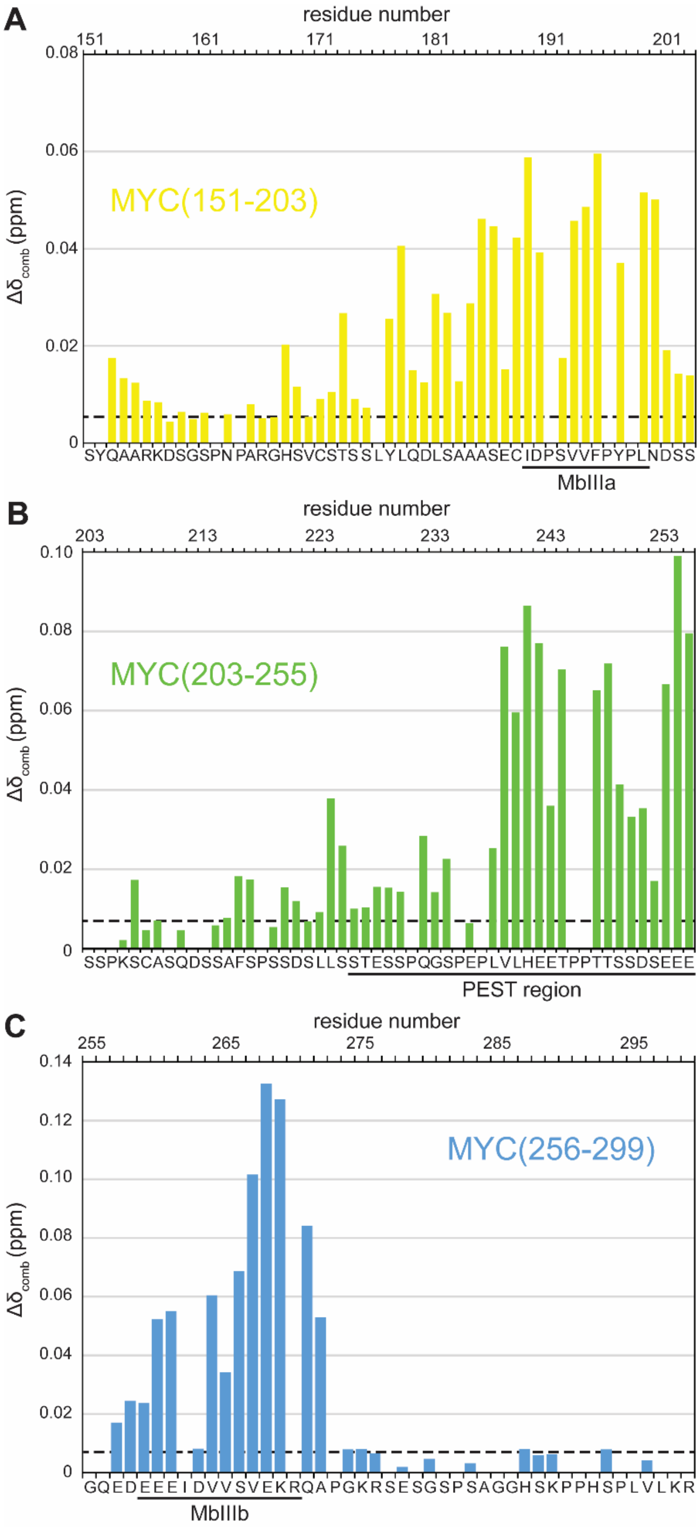


**Figure S10. Conserved motifs in the disordered MYC region bind to the MYC:MAX DBD.** (A-C) Quantitative CSP analysis for binding of the MYC:MAX DBD to the $^{15}$N-labeled MYC fragments 151–203 (A, yellow), 203–255 (B, green) and 256–299 (C, blue). The differences in chemical shift ($\Delta\delta_{comb}$) are calculated according to equation (1) and plotted against the amino acid sequence. The value above which CSPs are significant (dashed lines) is calculated as described in Materials and Methods.

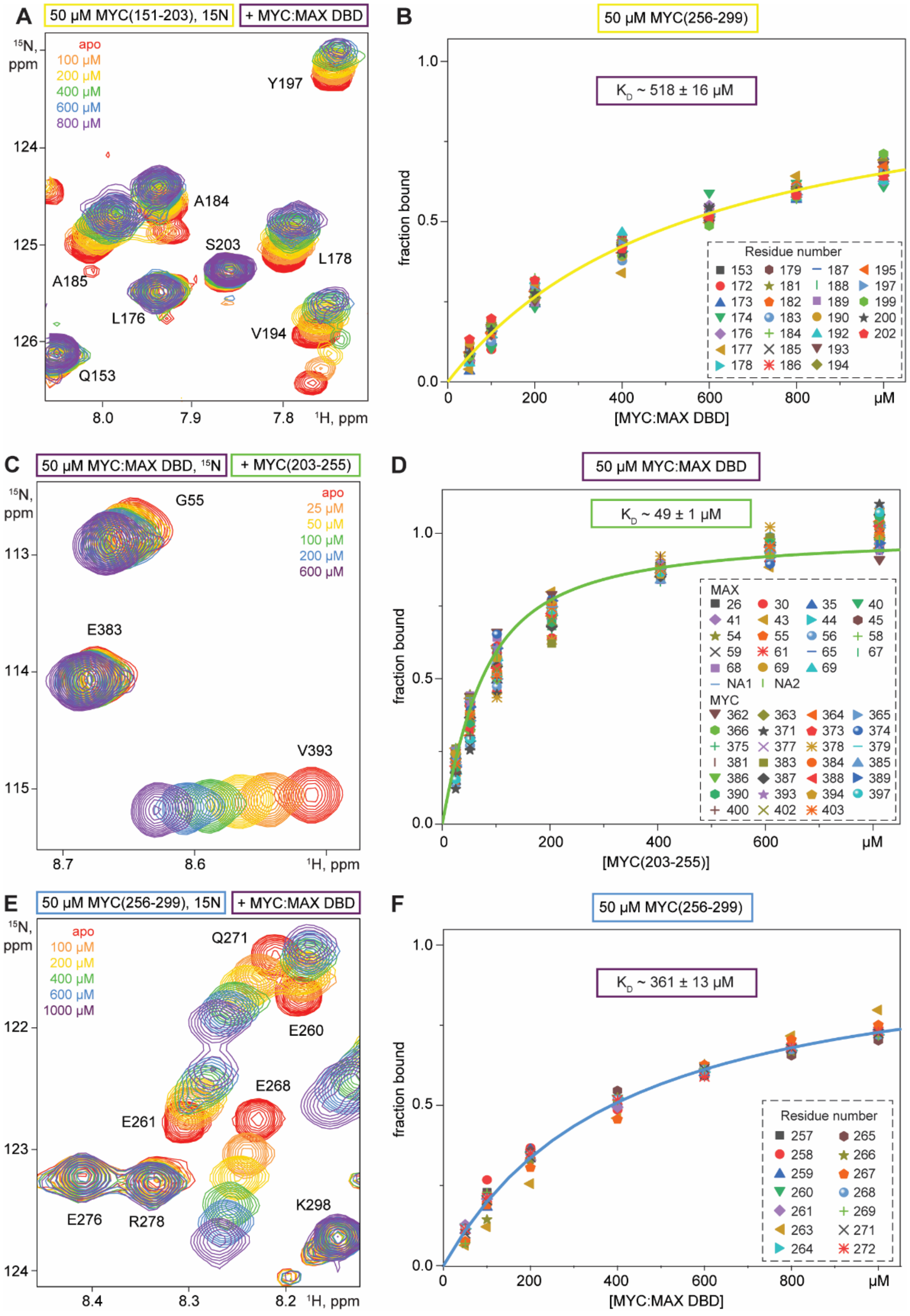


**Figure S11. Central MYC fragments bind the DBD with high micromolar affinity.** (A-C) Zoom on $^{1}$H-$^{15}$N HSQC spectra of 50 µM $^{15}$N-labeled (A) MYC(151–203), (B) MYC(203–255) and (C) MYC(265–299), in presence of increasing concentrations of MYC:MAX DBD, indicated by gradual coloring from red to purple. Assigned resonances are labeled. (D-F) Analysis of NMR titration data for the interactions shown in (A-C). For each of the indicated resonances, the fraction of bound protein was determined from the CSPs and plotted against the ligand concentration. The $K_D$ (± SD) is obtained from a global fit (solid lines) with equation (2).

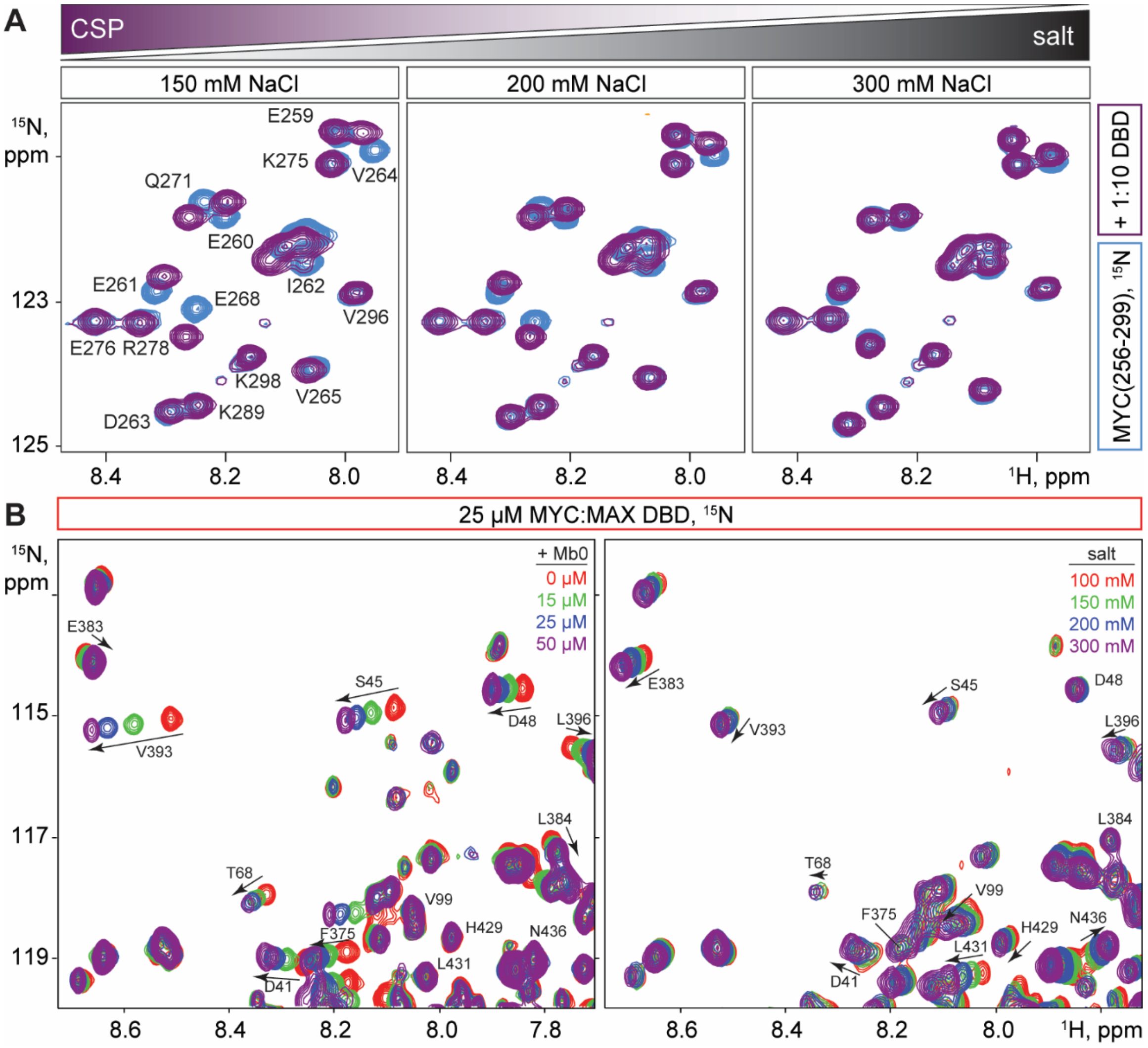


**Figure S12. Specific binding of disordered MYC fragments to the DBD is driven by electrostatic interactions.** (A) Zoom on $^{1}H$-$^{15}N$ HSQC spectra of 50 µM $^{15}N$-labeled MYC(256–299) in absence (blue) and presence of 500 µM MYC:MAX DBD (purple) in buffer containing 150, 200 or 300 mM NaCl (from left to right). CSPs become smaller with increasing salt concentrations. (B) $^{1}H$-$^{15}N$ TROSY spectra of 25 µM $^{15}N$-labeled MYC:MAX DBD in presence of increasing concentrations of Mb0 (left) or NaCl (right), indicated by gradual coloring from red to purple. The CSP pattern differs significantly with regards to directionality, extent, and affected residues. Assignments and CSPs are indicated for a selection of residues that are affected either by addition of Mb0 or NaCl.

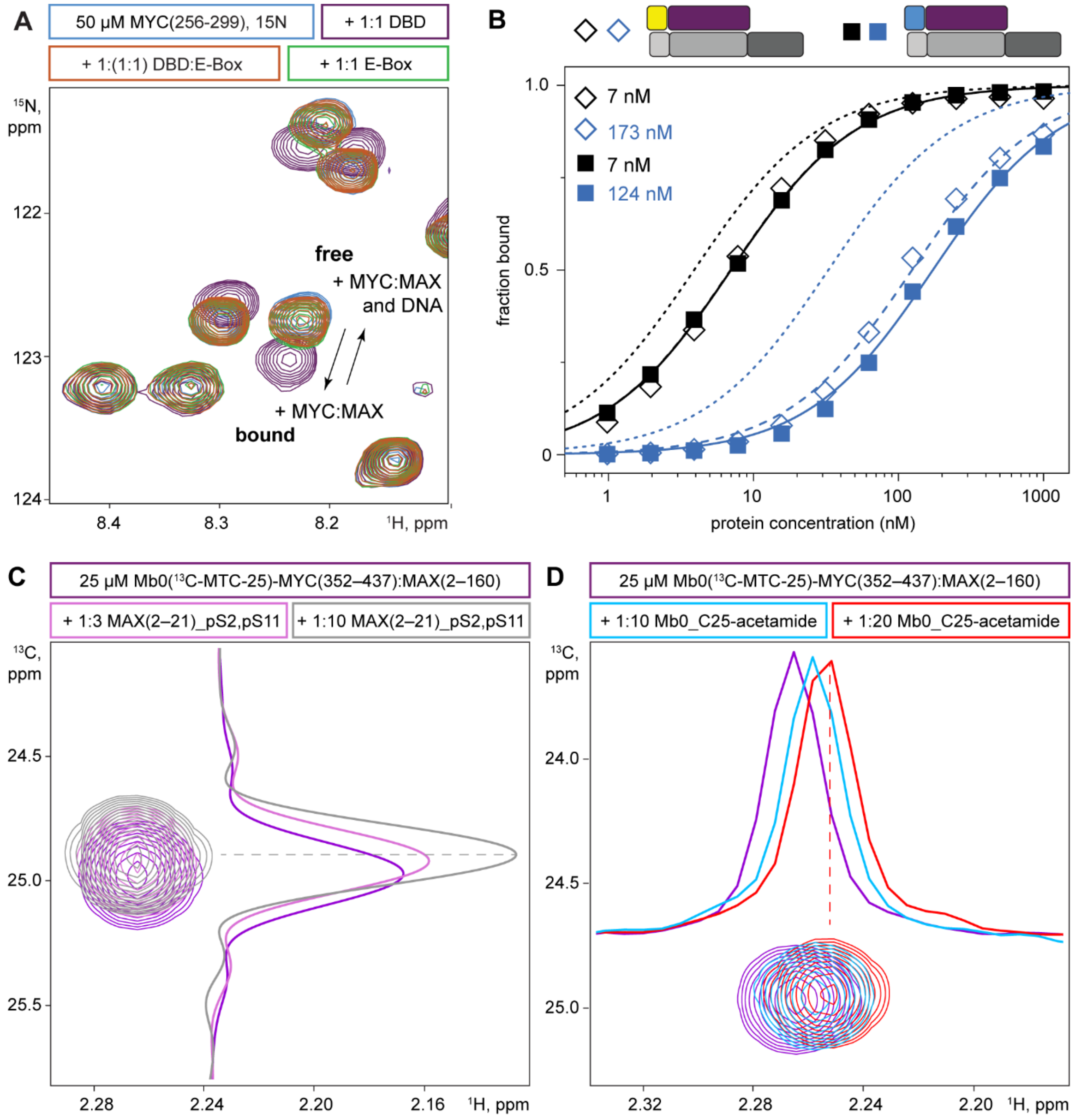


**Figure S13. MYC boxes and the MAX N-terminal peptide compete with DNA about the same binding site on the DBD.** (A) $^{1}$H-$^{15}$N HSQC spectra of 50 µM MYC(256–299) in its free form (blue) or in presence of equimolar amounts of E-Box DNA (green), MYC:MAX DBD (purple) or DBD and E-Box-DNA. (B) Steady-state analysis for the binding of MbIIIa-MYC(352–437):MAX(2–160) (diamonds, dashed lines) and MbIIIb-MYC(352–437):MAX(2–160) (squares, solid lines) to immobilized E-Box (black) and mutant E-Box 2 DNA (blue). $K_D$ values are obtained from a global fit with equation (3), see Materials and Methods, assuming a 1:1 binding model. For comparison, the dotted lines represent the respective fits for MYC(352–437):MAX(2–160) as in Figure4B. (C) $^{1}$H-$^{13}$C HMQC spectra of Mb0-MYC(352–437):MAX(2–160), that is $^{13}$C-MTC labeled on residue C25, in absence (purple) or in presence of a 3- (light pink) or 10-fold (gray) molar excess of pS2, pS11-double phosphorylated MAX N-terminal peptide. (D) as in (C), but in presence of a 10- (cyan) or 20-fold (red) molar excess of Mb0 peptide that is acetamide-labeled on C25 to prohibit disulfide shuffling. 1D cross-sections through the MTC-25 resonances in (C) and (D) illustrate the CSPs upon addition of free peptides.

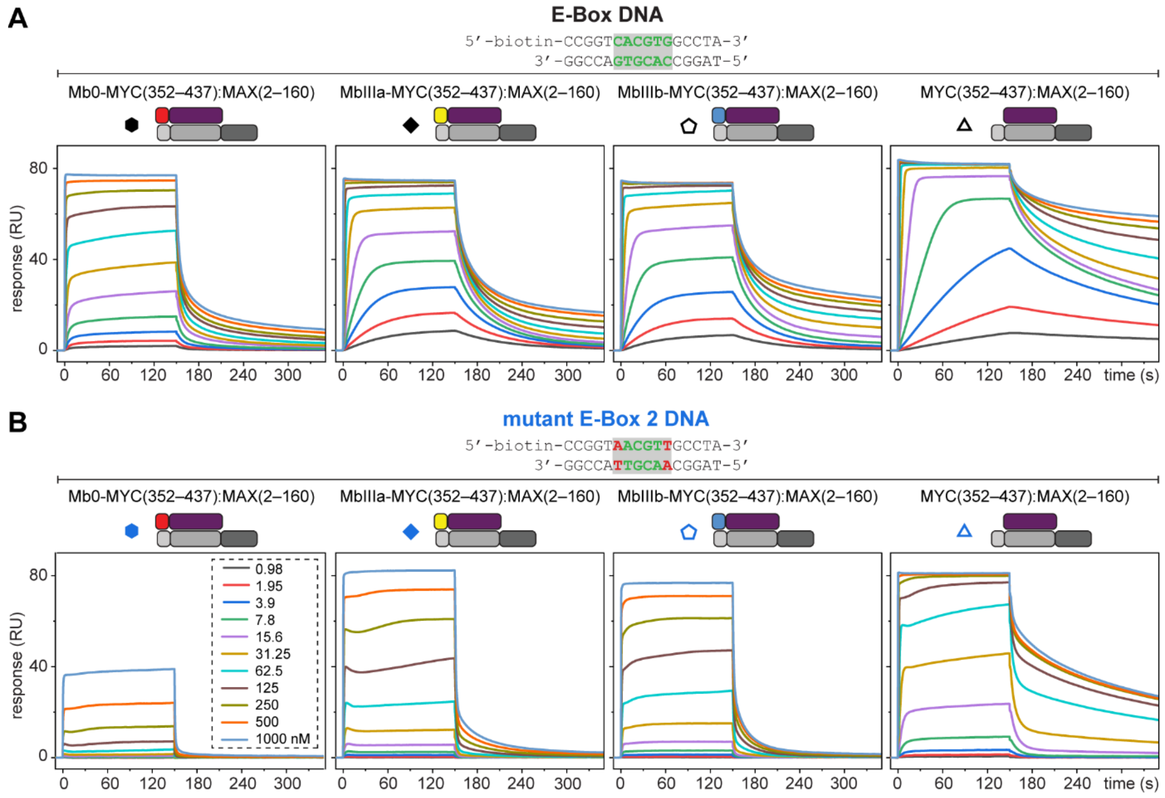


**Figure S14. MYC boxes that are directly tethered to the DBD reduce the affinity for DNA and accelerate dissociation**. (A) SPR sensorgrams for the binding of Mb0-, MbIIIa- and MbIIIb-linked and unmodified MYC(352–437):MAX(2–160) (from left to right) to immobilized E-Box DNA. (B) as in (A) but for binding to mutant E-Box 2 DNA. Altered nucleotides in the hexameric consensus sequence are highlighted in red.

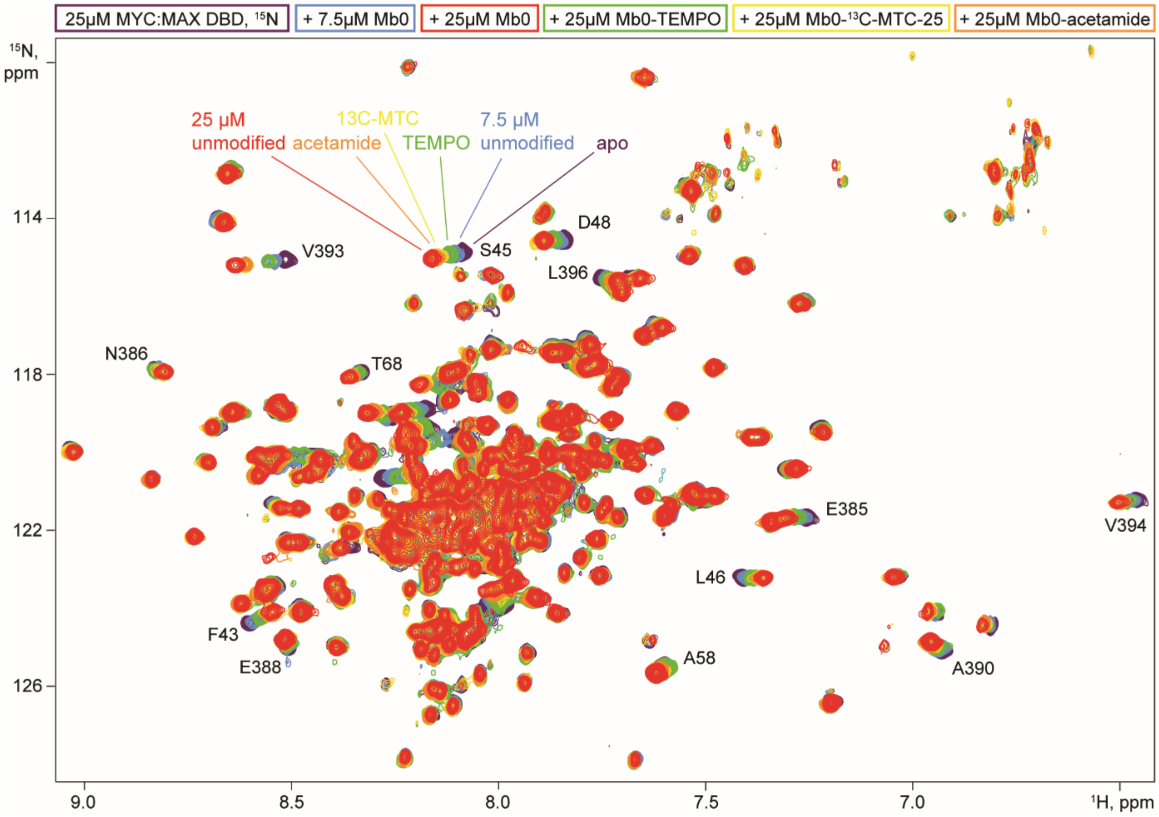


**Figure S15. Binding of Mb0 to the MYC:MAX DBD tolerates covalent modifications on C25.** $^{1}$H-$^{15}$N TROSY spectra of 25 µM MYC(352–437):MAX(22–102) in absence (purple) and presence of different Mb0 peptides: 25 µM unmodified Mb0 (red), 25 µM acetamide-labeled Mb0 (orange), 25 µM $^{13}$C-MTC-labeled Mb0 (yellow), 25 µM TEMPO-labeled Mb0 (green) and 7.5 µM unmodified Mb0 (blue). Assignments for some residues are indicated

**Table S1. DNA duplex oligonucleotides used in this study**. The 5’ and 3’ ends are indicated. The E-Box sequence is in green. Nucleobases that are modified in mutant E-Box sequences are in red.

| DNA | Sequence | Method |
|---|---|---|
| E-Box | 5'-biotin-CCGGT**CACGTG**GCCTA-3’<br>3'-GGCCA**GTGCAC**CGGAT-5’ | SPR |
| E-Box | 5'-CCGGT**CACGTG**GCCTA-3’<br>3'-GGCCA**GTGCAC**CGGAT-5’ | NMR |
| Mutant E-Box 1 | 5'-biotin-CCGGT**CATATG**GCCTA-3’<br>3'-GGCCA**GTATAC**CGGAT-5’ | SPR |
| Mutant E-Box 2 | 5’-biotin-CCGGT**AACGTT**GCCTA-3’<br>3’-GGCCA**TTGCAA**CGGAT-5’ | SPR |
| Negative control | 5’-biotin-CCGGT**TTAGCA**GCCTA-3’<br>3’-GGCCA**AATCGT**CGGAT-5’ | SPR |

**Table S2. Protein construct used in this study.** Protease cleavage sites are in orange, linker sequences in grey, solubility tags in green and affinity tags in blue. The target protein sequence is in black. The methionine residue from the translation start site in untagged MYC and MAX proteins is cleaved co-translationally.

| Protein (boundaries) | Purification tag / sequence |
|---|---|
| **MYC(1–88)** | N-His6-ZZ-TEV |
| | MKTHHHHHHGAQHDEAVDNKFNKEQQNAFYEILHLPNLNEEQRNAFIQSLKDDPSQSAN<br>LLAEAKKLNDAQAPKVDNKFNKEQQNAFYEILHLPNLNEEQRNAFIQSLKDDPSQSANLL<br>AEAKKLNDAA<br>PKVDANGGGSGGGSGGGSENLYFQGMPLNVSFTNRNYDLDYDSVQPYFYCDEEENFYQ<br>QQQQSELQPPAPSEDIWKKFELLPTPPLSPSRRSGLCSPSYVAVTPFSLRGDNDG |
| **MYC(89–150)** | N-His6-ZZ-TEV |
| | {tag see above} GGSFSTADQLEMVTELLGGDMVNQSFICDPDDETFIKNIIIQDCMWSG<br>FSAAAKLVSEKLA |
| **MYC(151–203)** | N-His6-ZZ-TEV |
| | {tag see above}<br>SYQAARKDSGSPNPARGHSVCSTSSLYLQDLSAAASECIDPSVVFPYPLNDSS |
| **MYC(151–255)** | N-His6-ZZ-TEV |
| | {tag see above}<br>SYQAARKDSGSPNPARGHSVCSTSSLYLQDLSAAASECIDPSVVFPYPLNDSSSPK<br>SCASQDSSAFSPSSDSLLSSTESSPQGSPEPLVLHEETPPTTSSDSEEE |
| **MYC(203–255)** | N-His6-ZZ-TEV |
| | {tag see above} SSPKSCASQDSSAFSPSSDSLLSSTESSPQGSPEPLVLHEETPPTTSSDSEEE |
| **MYC(256–299)** | N-His6-ZZ-TEV |
| | {tag see above} GQEDEEEIDVVSVEKRQAPGKRSESGSPSAGGHSKPPHSPLVLKR |
| **MYC(256–351)** | N-His6-ZZ-TEV |
| | {tag see above}<br>GQEDEEEIDVVSVEKRQAPGKRSESGSPSAGGHSKPPHSPLVLKRCHVSTHQHN<br>YAAPPSTRKDYPAAKRVKLDSVRVLRQISNNRKCTSPRSSDTE |
| **MYC(303–351)** | N-His6-ZZ-TEV |
| | {tag see above} STHQHNYAAPPSTRKDYPAAKRVKLDSVRVLRQISNNRKCTSPRSSDTE |
| **MYC(2–439)** | - |
| | MPLNVSFTNRNYDLDYDSVQPYFYCDEEENFYQQQQQSELQPPAPSEDIWKKFELLPTPP<br>LSPSRRSGLCSPSYVAVTPFSLRGDNDGGGGSFSTADQLEMVTELLGGDMVNQSFICDPD<br>DETFIKNIIIQDCMWSGFSAAAKLVSEKLASYQAARKDSGSPNPARGHSVCSTSSLYLQDL<br>SAAASECIDPSVVFPYPLNDSSSPKSCASQDSSAFSPSSDSLLSSTESSPQGSPEPLVLHEETP<br>PTTSSDSEEEQEDEEEIDVVSVEKRQAPGKRSESGSPSAGGHSKPPHSPLVLKRCHVSTHQ<br>HNYAAPPSTRKDYPAAKRVKLDSVRVLRQISNNRKCTSPRSSDTEENVKRRTHNVLERQR<br>RNELKRSFFALRDQIPELENNEKAPKVVILKKATAYILSVQAEEQKLISEEDLLRKRREQLK<br>HKLEQLRNSCA |
| **MYC(2–439)_ C0,C25**<br><br>**“Cys-light”** | - |
| | MPLNVSFTNRNYDLDYDSVQPYFYADEEENFYQQQQQSELQPPAPSEDIWKKFELLPTPP<br>LSPSRRSGLASPSYVAVTPFSLRGDNDGGGGSFSTADQLEMVTELLGGDMVNQSFIADPD<br>DETFIKNIIIQDAMWSGFSAAAKLVSEKLASYQAARKDSGSPNPARGHSVASTSSLYLQDL<br>SAAASEAIDPSVVFPYPLNDSSSPKSAASQDSSAFSPSSDSLLSSTESSPQGSPEPLVLHEETP<br>PTTSSDSEEEQEDEEEIDVVSVEKRQAPGKRSESGSPSAGGHSKPPHSPLVLKRAHVSTHQ<br>HNYAAPPSTRKDYPAAKRVKLDSVRVLRQISNNRKATSPRSSDTEENVKRRTHNVLERQR<br>RNELKRSFFALRDQIPELENNEKAPKVVILKKATAYILSVQAEEQKLISEEDLLRKRREQLK<br>HKLEQLRNSAA |

| | |
|---|---|
| **MYC(11–34)**<br><br>**“Mb0”** | N-His6-MBP-TEV<br>MKTHHHHHHGKIEEGKLVIWINGDKGYNGLAEVGKKFEKDTGIKVTVEHPDKLEEKFP<br>QVAATGDGPDIIFWAHDRFGGYAQSGLLAEITPDKAFQDKLYPFTWDAVRYNGKLIAY<br>PIAVEALSLIYNKDLLPNPPKTWEEIPALDKELKAKGKSALMFNLQEPYFTWPLIAADGG<br>YAFKYENGKYDIKDVGVDNAGAKAGLTFLVDLIKNKHMNADTDYSIAEAAFNKGETA<br>MTINGPWAWSNIDTSKVNYGVTVLPTFKGQPSKPFVGVLSAGINAASPNKELAKEFLEN<br>YLLTDEGLEAVNKDKPLGAVALKSYEEELAKDPRIAATMENAQKGEIMPNIPQMSAFW<br>YAVRTAVINAASGRQTVDEALKDAQTRITKGGGSGGGSGGGSLENLYFQGNYDLDYDS<br>VQPYFYCDEEENFYQQ |
| **MAX(2-160)** | N-Strep-TEV<br>MKTWSHPQFEKGSENLYFQSDNDDIEVESDEEQPRFQSAADKRAHHNALERKRRDHIK<br>DSFHSLRDSVPSLQGEKASRAQILDKATEYIQYMRRKNHTHQQDIDDLKRQNALLEQQ<br>VRALEKARSSAQLQTNYPSSDNSLYTNAKGSTISAFDGGSDSSSESEPEEPQSRKKLRME<br>AS |
| **MYC(352–437): MAX(22–102)** | N-His6 on MYC<br>MKTHHHHHHENVKRRTHNVLERQRRNELKRSFFALRDQIPELENNEKAPKVVILKKAT<br>AYILSVQAEEQKLISEEDLLRKRREQLKHKLEQLRNS<br><br>MADKRAHHNALERKRRDHIKDSFHSLRDSVPSLQGEKASRAQILDKATEYIQYMRRKN<br>HTHQQDIDDLKRQNALLEQQVRAL |
| **MYC(352–437): MAX(2–160)** | N-His6 on MYC<br>MKTHHHHHHENVKRRTHNVLERQRRNELKRSFFALRDQIPELENNEKAPKVVILKKAT<br>AYILSVQAEEQKLISEEDLLRKRREQLKHKLEQLRNS<br><br>MSDNDDIEVESDEEQPRFQSAADKRAHHNALERKRRDHIKDSFHSLRDSVPSLQGEKAS<br>RAQILDKATEYIQYMRRKNHTHQQDIDDLKRQNALLEQQVRALEKARSSAQLQTNYPS<br>SDNSLYTNAKGSTISAFDGGSDSSSESEPEEPQSRKKLRMEAS |
| **Mb0- MYC(352–437): MAX(2–160)** | N-His6-TEV on MYC(11–34)-MYC(352–437)<br>MKTHHHHHHGGGSGGGSGGGSENLYFQGNYDLDYDSVQPYFYCDEEENFYQQGGGSS<br>DNDDIEVESDEEQPRFQSAENVKRRTHNVLERQRRNELKRSFFALRDQIPELENNEKAP<br>KVVILKKATAYILSVQAEEQKLISEEDLLRKRREQLKHKLEQLRNS<br><br>MADKRAHHNALERKRRDHIKDSFHSLRDSVPSLQGEKASRAQILDKATEYIQYMRRKN<br>HTHQQDIDDLKRQNALLEQQVRAL |
| **MbIIIa- MYC(352–437): MAX(2–160)** | N-His6-TEV on MYC(175-200)-MYC(352–437)<br>MKTHHHHHHGGGSGGGSGGGSENLYFQSLYLQDLSAAASECIDPSVVFPYPLNGGGSS<br>DNDDIEVESDEEQPRFQSAENVKRRTHNVLERQRRNELKRSFFALRDQIPELENNEKAP<br>KVVILKKATAYILSVQAEEQKLISEEDLLRKRREQLKHKLEQLRNS<br><br>MADKRAHHNALERKRRDHIKDSFHSLRDSVPSLQGEKASRAQILDKATEYIQYMRRKN<br>HTHQQDIDDLKRQNALLEQQVRAL |
| **MbIIIb- MYC(352–437): MAX(2–160)** | N-His6-TEV on MYC(256-273)-MYC(352–437)<br>MKTHHHHHHGGGSGGGSGGGSENLYFQGQEDEEEIDVVSVEKRQAPGGGSSDNDDIEV<br>ESDEEQPRFQSAENVKRRTHNVLERQRRNELKRSFFALRDQIPELENNEKAPKVVILKK<br>ATAYILSVQAEEQKLISEEDLLRKRREQLKHKLEQLRNS<br><br>MADKRAHHNALERKRRDHIKDSFHSLRDSVPSLQGEKASRAQILDKATEYIQYMRRKN<br>HTHQQDIDDLKRQNALLEQQVRAL |
| **MYC(151–439): MAX(2–160)** | N-His6-Thrombin-Sumo on MYC; N-Flag-TEV on MAX<br>MGSSHHHHHHSSGLVPRGSHMASMSDSEVNQEAKPEVKPEVKPETHINLKVSDGSSEIF<br>FKIKKTTPLRRLMEAFAKRQGKEMDSLRFLYDGIRIQADQTPEDLDMEDNDIIEAHREQI<br>GGSMSYQAARKDSGSPNPARGHSVCSTSSLYLQDLSAAASECIDPSVVFPYPLNDSSSPK<br>SCASQDSSAFSPSSDSLLSSTESSPQGSPEPLVLHEETPPTTSSDSEEEQEDEEEIDVVSVEK<br>RQAPGKRSESGSPSAGGHSKPPHSPLVLKRCHVSTHQHNYAAPPSTRKDYPAAKRVKL<br>DSVRVLRQISNNRKCTSPRSSDTEENVKRRTHNVLERQRRNELKRSFFALRDQIPELENN<br>EKAPKVVILKKATAYILSVQAEEQKLISEEDLLRKRREQLKHKLEQLRNSCA<br><br>MDYKDDDDKGSENLYFQSDNDDIEVESDEEQPRFQSAADKRAHHNALERKRRDHIKDS<br>FHSLRDSVPSLQGEKASRAQILDKATEYIQYMRRKNHTHQQDIDDLKRQNALLEQQVR<br>ALKARSSAQLQTNYPSSDNSLYTNAKGSTISAFDGGSDSSSESEPEEPQSRKKLRMEAS |
